# Ultrasound stimulation of the motor cortex during tonic muscle contraction

**DOI:** 10.1101/2021.05.03.442483

**Authors:** Ian S. Heimbuch, Tiffany Fan, Allan Wu, Guido C. Faas, Andrew C. Charles, Marco Iacoboni

## Abstract

Transcranial ultrasound stimulation (tUS) shows potential as a noninvasive brain stimulation (NIBS) technique, offering increased spatial precision compared to other NIBS techniques. However, its reported effects on primary motor cortex (M1) are limited. We aimed to better understand tUS effects in human M1 by performing tUS of the hand area of M1 (M1_hand_) during tonic muscle contraction of the index finger. Stimulation during muscle contraction was chosen because of the transcranial magnetic stimulation-induced phenomenon known as cortical silent period (cSP), in which transcranial magnetic stimulation (TMS) of M1_hand_ involuntarily suppresses voluntary motor activity. Since cSP is widely considered an inhibitory phenomenon, it presents an ideal parallel for tUS, which has often been proposed to preferentially influence inhibitory interneurons. Recording electromyography (EMG) of the first dorsal interosseous (FDI) muscle, we investigated effects on muscle activity both during and after tUS. We found no change in FDI EMG activity concurrent with tUS stimulation. Using single-pulse TMS, we found no difference in M1 excitability before versus after sparsely repetitive tUS exposure. Using acoustic simulations in models made from structural MRI of the participants that matched the experimental setups, we estimated in-brain pressures and generated an estimate of cumulative tUS exposure experienced by M1_hand_ for each subject. We were unable to find any correlation between cumulative M1_hand_ exposure and M1 excitability change. We also present data that suggest a TMS-induced MEP always preceded a near-threshold cSP.

## Introduction

Transcranial ultrasound stimulation (tUS) has gained attention in the past years as a potential new tool for noninvasive brain stimulation (NIBS). tUS has higher spatial precision compared to other NIBS techniques such as transcranial magnetic stimulation (TMS) and transcranial electric stimulation (TES), which presents a possibility of improved targeting (Deffieux et al., 2013; Lee et al., 2015; Legon et al., 2014; Roth, Amir, Levkovitz, & Zangen, 2007; Wagner, Valero-Cabre, & Pascual-Leone, 2007). Furthermore, tUS can deliver its energy much deeper while maintaining focal precision—deeper than TMS or TES (Legon, Ai, Bansal, & Mueller, 2018).

Previous studies have demonstrated that ultrasound is capable of stimulating central structures in animals (Fry, Ades, & Fry, 1958), peripheral nerve pathways in animals and humans (Gavrilov et al., 1976; Gavrilov, Tsirulnikov, & Davies, 1996), the retina (Menz, Oralkan, Khuri-Yakub, & Baccus, 2013), and intact brain circuits in animals (Tufail et al., 2010). However, the number of human tUS studies thus far is limited. In primary somatosensory cortex, tUS has been shown to modulate touch discrimination (Legon et al., 2014), induce localized somatosensations when targeting the cortical hand representation (Lee et al., 2015, 2017), and induce changes in intrinsic and evoked EEG dynamics (Mueller, Legon, Opitz, Sato, & Tyler, 2014). In primary visual cortex, tUS can induce individual visual phosphenes, percepts of a flash of light, that were accompanied with an evoked potential and blood-oxygenation-level-dependent (BOLD) contrast similar to those seen with photic stimulation (Lee et al., 2016). However, the effects of tUS in primary motor cortex (M1) have been less clear.

Much of our understanding of motor cortex stimulation comes from TMS investigations, where it has been the standard for noninvasive M1 perturbation for decades (Chen et al., 1997; Hess, Mills, & Murray, 1987; Hess, Mills, Murray, & Schriefer, 1987; Huang, Edwards, Rounis, Bhatia, & Rothwell, 2005; Oberman, Edwards, Eldaief, & Pascual-Leone, 2011; Priori, Berardelli, Rona, Accornero, & Manfredi, 1998). Suprathreshold single-pulse TMS of M1 induces contraction in the corresponding muscles, and electromyography (EMG) allows for the quantification of these motor evoked potentials (MEPs). Since MEP size increases as a sigmoidal function of TMS intensity above motor threshold (Hess, Mills, & Murray, 1987; Möller, Arai, Lücke, & Ziemann, 2009), MEP strength is frequently used as an indicator of corticospinal excitability, both in neuromodulatory and behavioral interventions (Chen et al., 1997; Christov-Moore, Sugiyama, Grigaityte, & Iacoboni, 2016; Fitzgerald, Fountain, & Daskalakis, 2006; Hess, Mills, & Murray, 1987; Hess, Mills, Murray, et al., 1987; Huang et al., 2005; Liebetanz et al., 2003; Oberman et al., 2011; Priori et al., 1998; Thut & Miniussi, 2009). This is supported by pharmacological evidence that shows motor threshold, the TMS intensity needed to elicit an MEP, is a proxy for the within-subject excitability of the cortico-cortical axons affected by the induced current of TMS pulse (Ziemann et al., 2015). We investigated the neuromodulatory effects of tUS on M1 by analyzing its effect on TMS-evoked MEP.

Cortical silent periods (cSP) are a phenomenon of suppressed EMG activity during tonic contraction following single-pulse TMS of the corresponding M1 motor representation. cSPs typically last from 100-300 ms. Importantly, cSPs are considered to be driven predominantly by cortical inhibition from ~50 ms after instigation (Wolters, Ziemann, & Benecke, 2012). Specifically, pharmacological evidence suggests that the cSP effect is mediated by GABA receptor-dependent postsynaptic inhibition (Werhahn, Kunesch, Noachtar, Benecke, & Classen, 1999; Ziemann et al., 2015). When elicited during tonic contraction of the contralateral hand, cSPs are reported to be observable either following an MEP or without inducing an MEP, at subthreshold TMS intensities (Classen & Benecke, 1995; Davey, Romaiguère, Maskill, & Ellaway, 1994). As such, cSP provides a valuable method of investigation of inhibitory mechanism in motor cortex.

As a whole, the field is still building its understanding of what, if anything, tUS can affect via M1 stimulation. To date, no tUS study has been able to induce motor contraction through human M1 stimulation. Previous animal model studies suggest that tUS-induced MEPs may only be producible at acoustic intensities above human-safe levels (Kim, Chiu, Lee, Fischer, & Yoo, 2014; King, Brown, Newsome, & Pauly, 2013; Krasovitski, Frenkel, Shoham, & Kimmel, 2011; Tufail et al., 2010; Ye, Brown, & Pauly, 2017; Yoo et al., 2011). Given that change in motor contraction strength has been the benchmark for M1 modulation studies, the established capabilities of TMS have been leveraged to investigate tUS effects on M1. For example, tUS of the hand area reduced the size of motor evoked potentials (MEPs) evoked by concurrent and concentric TMS (Legon, Bansal, Tyshynsky, Ai, & Mueller, 2018). A separate study reported a lasting increase in the size of MEPs after exposure to an ultrasound imaging device (Gibson et al., 2018). Additionally, tUS of M1 alone was shown to affect reaction time in a motor task (Legon, Bansal, et al., 2018).

Because of its physiological underpinnings, the cSP is in a unique position to be leveraged as an externally detectable phenomenon to better understand tUS effects on M1. Specifically, tUS has been proposed to preferentially affect inhibitory interneurons (Kim et al., 2014; Legon et al., 2014; Nguyen, Berisha, Konofagou, & Dmochowski, 2020; Plaksin, Kimmel, & Shoham, 2016; Rinaldi, Jones, Reines, & Price, 1991), feeding well into cSP’s existence as a interneuron-facilitated phenomenon. Additionally, since cSPs have been reported to occur without a preceding MEP (Cantello, Gianelli, Civardi, & Mutani, 1992; Classen & Benecke, 1995; Davey et al., 1994; Wassermann et al., 1993; Wolters et al., 2012), tUS’ apparent inability to instigate an MEP does not preclude its use to attempt induction of cSPs. But as of yet, it is unknown if tUS can engage the necessary inhibitory circuits to instigate a cSP. We addressed this by performing tUS of M1 on participants executing voluntary muscle contraction and analyzing the EMG data from the contracted muscle.

## Methods

Two experiments were performed in this study. In Experiment 1, we measured how tonically contracted hand muscles respond to single-pulse TMS and single-burst tUS of M1. We performed separate trials using tUS and TMS. In Experiment 2, we measured how cortical excitability is affected by tUS exposure. Excitability was gauged using single-pulse TMS.

### Data acquisition

#### Participant Demographics

Research participants were right-handed with no neurological conditions. Due to TMS use, subjects with an increased risk of seizure were excluded (Supplemental Figure 14). Due to MRI use, subjects with MR-incompatible implants were excluded. Participants were 18 to 42 years old, A subset of Experiment 1 participants (n = 10; mean: 25.9 years) participated in Experiment 2 (n = 8; mean: 26.75 years). Note that subject ID numbers are not sequential since other recruited subjects were used in a different study.

#### EMG and NIBS Placement

Electrode sites were cleaned with abrasive skin prep gel (Nuprep) and alcohol wipes. A surface EMG electrode (two 10 × 1 mm contacts; 10 mm spacing) measured the right first dorsal interosseous (FDI) muscle activity, and the signal was amplified (x1000) (Delsys Inc., Boston, MA) and sampled at 5000 Hz. The surface electrode was additionally secured to the finger with medical tape. A wide ground electrode was placed on the back of the hand. EMG was recorded for 1-second epochs around NIBS (both TMS and tUS) onset. A structural MRI (T1-weighted; 0.8 × 0.8 × 0.8 mm voxels) was acquired in a previous visit, and each structural MRI was registered to standard space for NIBS targeting of standard-space coordinates (Montreal Neurological Institute, MNI; 1-mm atlas). Registration was performed in FSL (the FMRIB Software Library) on brain volumes extracted using the optiBET tool for FSL’s BET (Jenkinson, Beckmann, Behrens, Woolrich, & Smith, 2012; Lutkenhoff et al., 2014). NIBS position with respect to the subject’s head was tracked using neuronavigation software (Brainsight, Rogue Research, Montreal, QC) loaded with the subject’s MRI. The neuronavigation software was prepared with pre-determined trajectories, which were the shortest Euclidean distance from the scalp to a set voxel as determined by a custom MATLAB script (Mathworks, Inc., Natick, MA). At the beginning of each NIBS session, five single TMS pulses were given at each target of a 6-target grid over the left motor cortex using MNI space (Supplemental Figure 12). The grid’s origin was placed at MNI coordinates that correspond to M1_hand_ as based on a meta-analysis of fMRI motor experiments: x = −39, y = −24, z = 57 (Mayka, Corcos, Leurgans, & Vaillancourt, 2006). This coordinate corresponds morphologically to the cortical ‘hand knob’ (Yousry et al., 1997). The other five targets on the grid were in a 12 voxel-width grid (9.6 mm grid interval) anterior, posterior, and medial, anteromedial, and posteromedial from the M1_hand_ coordinate in subject space. The targets that elicited the largest, second-largest, and third-largest average MEPs were used as placement points for the NIBS devices. These three positions are referred to below as “TMS target”, “2^nd^-best”, and “3^rd^-best” targets respectively. For all TMS trials, the TMS coil was oriented with the handle pointed backwards and angled 45° from midline.

#### Experiment 1, cSPs

Participants moved their index finger laterally during trials to maintain consistent FDI contraction across trials, as monitored by a digital scale (20-40% maximum voluntary contraction, depending on the subject). NIBS was delivered during contraction. Percent maximum contraction varied between subjects so that every subject maintained a comparable level of EMG activity (~150-200 peaks per second). 20 trials were performed at each of three TMS intensity levels: 90%, 100%, and 110% of % active motor threshold (aMT) (60 TMS trials total per subject). 20 tUS trials were performed for each of the following four parameters: 300-ms burst duration at the TMS target, 300-ms burst duration at the 2^nd^-best target, 300-ms burst duration at the 3^rd^-best target, and 500-ms burst duration at the TMS target. Subjects were told to use the feedback of the digital scale display to maintain their target FDI contraction force, and subjects were cued to relax between trials to avoid fatigue. Subjects monitoring their contraction level also had the benefit of keeping subject attention constant across trials in Experiment 1, since attention can affect EMG measurements (Vance, Wulf, Töllner, McNevin, & Mercer, 2004; Zachry, Wulf, Mercer, & Bezodis, 2005). Trials had a jittered intertrial interval of 10 ± 2 s.

#### Experiment 2, Cortical Excitability

MEPs were measured with the subject’s hand relaxed. TMS was delivered at the same suprathreshold intensity for both “before” and “after” conditions within subjects. TMS was set to 110-120 %rMT (percent resting motor threshold), (110%: n = 1; 115%: n = 4; 120% n = 3). %rMT was varied across subjects to assure that each subject had consistent MEP sizes. 20 MEPs were acquired before exposure to tUS and 20 MEPs were acquired after exposure to tUS. tUS exposure protocol was the same as described in “Experiment 1, cSPs:” (20 trials each: 300 ms at TMS target, 300 ms at 2^nd^-best, 300 ms at 3^rd^-best, 500 ms at TMS target). Trials had a jittered intertrial interval of 10 ± 2 s.

#### tUS Equipment

The tUS device used was a 500-kHz focused piezoelectric transducer (Blatek Industries, Inc., State College, PA). The transducer had a face width of 3 cm and a focal point of 3 cm. The transducer was housed in a custom 3D-printed handle, and an infrared tracker was mounted to the housing for neuronavigation (Figure 1). The transducer was driven by 500-kHz sine-wave voltage pulses from a waveform generator (33500B Series, Keysight Technologies, Santa Rosa, CA) and voltage pulses were amplified by a 50-dB radio frequency amplifier (Model 5048, Ophir RF, Los Angeles, CA). A 3-dB fixed attenuator was attached in line following the amplifier.

**Figure 1.**
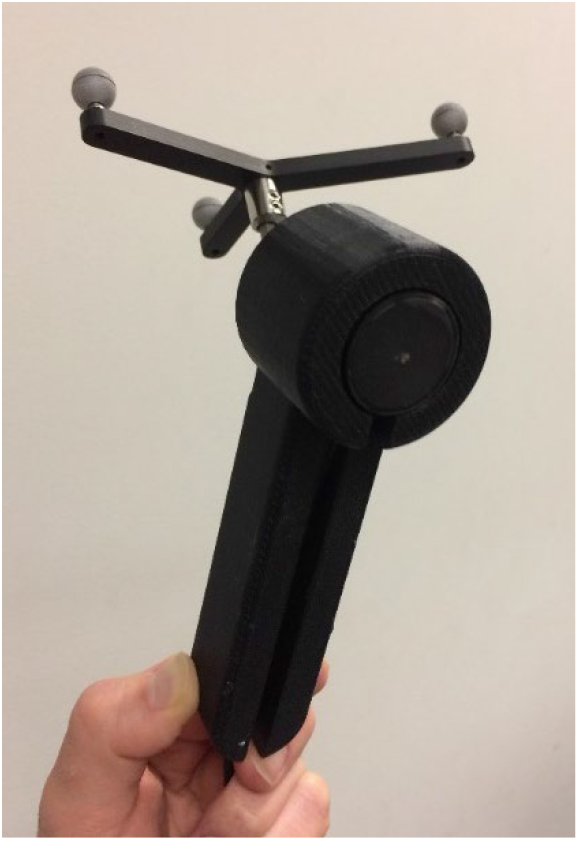
tUS transducer in housing

tUS bursts were pulsed with a 1-kHz pulse repetition frequency and a duty cycle of 36% (Figure 2). Each burst lasted either 300 or 500 ms (tUS on for 108 or 180 ms total). Transducer output was confirmed via measurements made via hydrophone in degassed water (1 mm, Precision Acoustics Ltd, Dorchester, UK). Transducer output was set to produce an intensity of 15.48 W/cm^2^ in degassed water (spatial peak, pulse average; I_sppa_). These parameters were chosen to not exceed an in-tissue estimate of 4.9 W/cm^2^. This is within safe levels (max Mechanical Index: 0.8) (ter Haar et al., 2011) and is within intensities of previous human tUS studies (Ai, Mueller, Grant, Eryaman, & Legon, 2016; Lee et al., 2015, 2016; Legon et al., 2014; Mueller et al., 2014).

**Figure 2.**
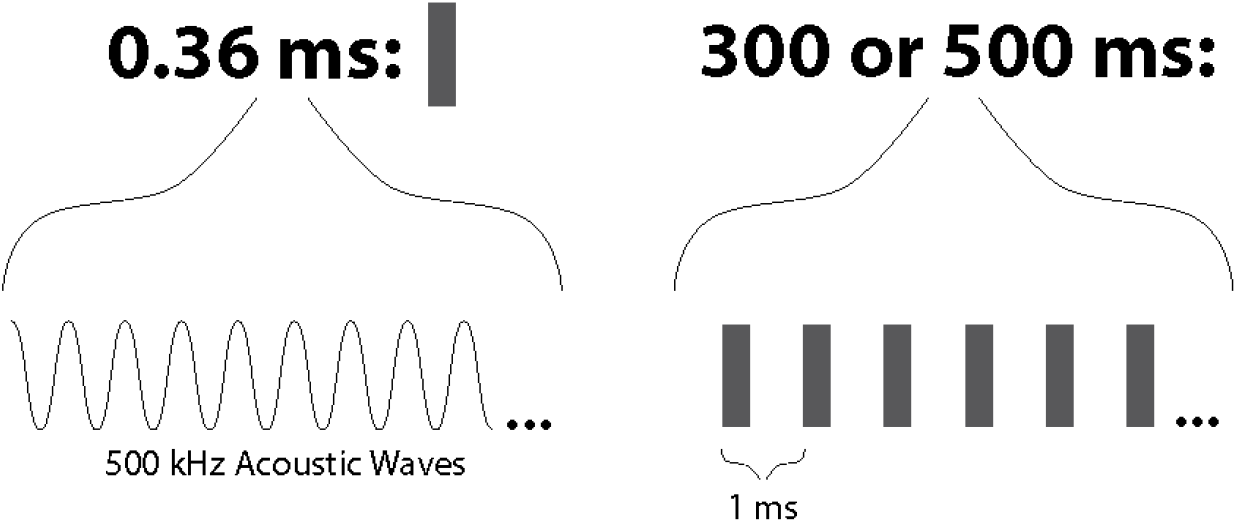
tUS Protocol. Illustration of a single trial of tUS.

#### TMS Equipment

Single-trial, monophasic TMS was applied using a figure-eight coil (70 mm diameter) via a Magstim 200^2^ magnetic stimulator (Magstim, Whitland, Dyfed, UK). An infrared tracker was mounted to the TMS coil for neuronavigation. Individual resting and active motor thresholds were determined using simple adaptive PEST (SA-PEST) (Adaptive PEST TMS threshold assessment tool, Brain stimulation laboratory, Department of Psychiatry, Medical University of South Carolina). Trials had a jittered intertrial interval of 10 ± 2 s.

### EMG data analysis

#### EMG Post-processing

EMG traces were high-pass filtered with a 10-Hz cutoff (filter transition: 5-10 Hz), unless noted as unfiltered. This high-pass filter was applied to remove voltage shift and low-frequency noise. All EMG post-processing was performed using MATLAB.

#### cSP Measurement

cSPs were measured using an automated script written in MATLAB, which used a rolling standard deviation (STD) to see when EMG activity quieted below a threshold (Supplemental Figure 1). The rolling STD window had a width of 3 ms. The cSP threshold was set using the baseline EMG variability—specifically ½ STD of the rolling STD trace the 200-ms period before TMS onset. cSP onset was set to the timepoint the rolling STD first fell below threshold after the MEP peak. cSP starts were also contingent on the raw EMG being near or below zero (specifically, below the same threshold value). In rare cases in which no cSP onset point was found within the first 15 ms, the onset was set to the first rising EMG value.

cSP offset was set where the rolling STD first rose back above threshold. A 15-ms ‘amnesty period’ was included for offset auto-detection to correct for trials with large MEPs, which have large STD values in their valleys after the MEP peak. If the rolling STD reached threshold from 0-15 ms after cSP onset, the window from 15-30 ms after cSP onset would be checked (‘post-amnesty period’). If the rolling STD trace stayed below threshold during the entire 15-ms post-amnesty period, the amnesty period transgression would be ignored (i.e. a silent period >30 ms). If not, the first threshold breach is used (i.e. a silent period <15 ms).

#### MEP Measurement, Resting

To quantify the size of MEPs Experiment 2, we used the area of the MEPs. The area under the curve (AUC) of the rectified MEP waveform from 20 to 120 ms after the TMS pulse was estimated via the trapezoidal method in MATLAB. AUC was not used in Experiment 1 (voluntary contractions) because of surrounding EMG activity.

#### MEP Measurement, Voluntary Contraction

MEP peak-to-peak measurements were measured by the absolute height from the peak to the mean of the two flanking valleys of the peak. To improve accuracy of automated MEP detection during tonic contraction, MEP search was constrained using per-subject exemplar data from trials with overt MEPs. A 10-ms search window was centered around the expected MEP timepoint. Expected MEP timepoint was the median MEP timepoint during resting TMS MEP trials (Experiment 2 TMS MEP data). For the two subjects who did not participate in Experiment 2, trials with visually overt MEPs during tonic contraction were used instead (Experiment 1 TMS MEP data). This search approach was used both for the positive MEP peak and its two flanking negative valleys.

Candidate peaks and valleys in the EMG data were found using the *findpeaks* MATLAB function. To avoid minor, extraneous peaks from being selected, only peaks with a prominence and width above the 50^th^ percentile were eligible.

TMS cSP trials were categorized into three labels: *MEP*, *stub*, and *none*. Overt MEPs (*MEP*) occurred within the 10-ms search window and had a peak-to-peak height above 0.5 mV. Potential-but-short MEPs (*stubs*) occurred within the 10-ms search window and had a peak-to-peak height below 0.5 mV. Trials with no peak that met conditions (50^th^ percentile prominence, width) within the 10-ms search window were labeled *none*.

#### EMG Characteristics

Additional characteristics of the EMG traces were calculated to contrast TMS and tUS effects on the FDI EMG signal during voluntary contraction, as well compare within different periods of tUS trials. First, spectral components were determined by estimating the short-term, time-localized power spectrum of each trial and then taking the mean to get separate average spectrograms for TMS trials and tUS trials. Second, lengths of silences in the EMG signal were calculated with a sliding window approach. Specifically, the cSP algorithm (see <u>cSP Measurement</u>) searched for a silence duration from a window centered at 0.001-second intervals from 0.05 to 0.95 s. The first and last 0.05 s were excluded to avoid edge artifacts. The results were then averaged within their respective groups to get mean silence traces.

Two additional characteristics were calculated to investigate possible EMG responses time-locked to tUS exposure: the height of the EMG (AUC) and the rate of EMG peaks. AUC was calculated as described above (MEP Measurement, Resting) for two 150-ms epochs: from 200 to 50 ms before tUS onset and from 50 to 200 ms after tUS onset. This provides two 150-ms epochs wholly covered by ‘off’ and ‘on’ periods of tUS. Rate of EMG peaks was calculated using findpeaks() function in MATLAB on each EMG trace, binning peaks by time for the tUS-off or tUS-on periods (i.e. both the pre- and post-tUS periods were included together for tUS-off). All EMG characteristics processing was performed in MATLAB.

### Acoustic simulation

#### Skull Mask Processing

For acoustic simulations and skull thickness measurements, binary skull masks were produced in BrainSuite using its “Cortical Surface Extraction Sequence” (Shattuck & Leahy, 2002). Skull masks were corrected by hand with the mask brush tool in BrainSuite. In MATLAB, skull masks were linearly interpolated to increase resolution to 0.2-mm-width voxels, and they were rotated such that the tUS trajectory was in line with the computational grid. Masks were also smoothed via morphological image processing both before and after transformation. Masks were cropped to the area of interest, creating a 484 × 484 × 484 volume.

#### k-Wave Simulations

Acoustic simulations were performed using k-Wave, an open-source acoustics toolbox for MATLAB (Treeby & Cox, 2010). Each skull mask was imported into k-Wave, providing a computational grid spacing of 0.2 mm. To simulate the transducer, we set a curved disc pressure source (k-Wave function: *makeBowl*) with a curvature radius of 30 mm and aperture of 30 mm to mirror the focal length and width of the real transducer, respectively. The pressure source emitted a 0.5 MHz sine wave, resulting in a grid points per wavelength (PPW) of 14.8. Simulations were performed at a temporal interval of 285 temporal points per period (PPP) for a Courant-Friedreichs-Lewy (CFL) number of 0.0519. Perfectly matched layers (PML) of 14 grid points were added for a total grid size of 512 × 512 × 512.

To allow comparison to real-world pressure measurements in the water tank, each tUS trajectory was simulated twice: once to simulate propagation through the skull and once to simulate propagation through water. For skull simulations, the same acoustic properties were given for all points within the skull mask: density of 1732 kg/m^3^, a sound speed of 2850 m/s, and an alpha coefficient of 8.83 [dB/(MHz^y^ cm)] (Treeby & Cox, 2014). The use of homogenous skull acoustic properties has been shown to be effective in simulations within the frequencies used here (Jones & Hynynen, 2016; Miller, Eames, Snell, & Aubry, 2015; Robertson, Cox, Jaros, & Treeby, 2017). All values not within the skull mask were given bulk acoustic values of brain: 1546.3 kg/m^3^, a sound speed of 1035 m/s, and an alpha coefficient of 0.646 [dB/(MHz^y^ cm)] (Duck, 1990). Homogenous water simulations were given acoustic properties of water at 20 °C: a density of 998 kg/m^3^, a sound speed of 1482 m/s, and an attenuation constant of 2.88 × 10^−4^ [Np / m] (Duck, 1990). An alpha power of 1.43 was used for all simulations.

To estimate in-brain pressures experienced by participants for a given tUS trajectory, we used a ratio of pressures from the skull and water simulations. The estimated pressure (*P_est_*.) at a given location was calculated as

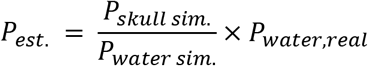

Where *P_skull sim_*. is the temporal maximum pressure value at that same location in the skull simulation for the specific subject and trajectory. *P_water sim_*. is the temporal maximum pressure value at the focal point of the water simulation. *P_water,real_* is the temporal maximum pressure value at the focal point measured in a water tank of degassed water using the same parameters as used in the experiment (see <u>tUS Equipment</u>). To avoid any potential outliers in the simulated data, spatial averaging was performed on *P_skull sim_*. and *P_water sim_*. by taking the mean within a 0.6-mm radius sphere. *P_water,real_* was 1.40 MPa for all subjects except one (sbj11), whose *P_water,real_* was 1.13 MPa due to the lower waveform generator setting used for that session (user error).

Simulations were performed on the Ahmanson-Lovelace Brain Mapping Center computational cluster. Each simulation instance was allocated 24 CPU cores and took approximately 2.5 hours with the C++ implementation of k-Wave (kspaceFirstOrder3D-OMP) (Treeby, Jaros, Rendell, & Cox, 2012).

#### Target Registration

NIBS targets and the location of M1_hand_ were determined via registration to standardized stereotactic space (Montreal Neurological Institute, MNI). Registration was performed with FSL’s FNIRT/FLIRT tools (Jenkinson, Bannister, Brady, & Smith, 2002; Jenkinson & Smith, 2001). M1_hand_ was set to the voxel closest to the MNI coordinates x = −39, y = −24, z = 57 (Mayka et al., 2006).

#### Exposure

An estimate of cumulative M1_hand_ exposure was made by multiplying the individual peak pressure at the M1_hand_ voxel for each of the three tUS trajectories by the time the tUS device was on for that location. Specifically, exposure was defined as

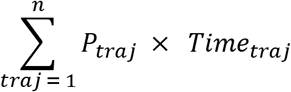

for ***n*** number of tUS trajectories, where ***P_traj_*** is the pressure at the M1_hand_ voxel for that trajectory, and ***Time_traj_*** is time tUS was on for that trajectory. We display these values in the form Pascal-hours (Pa·hr).

### Statistics

#### Experiment 1, cSPs

TMS cSP durations vs. aMT was analyzed with a one-way repeated-measures ANOVA. For post-hoc tests, Welch’s t-tests were performed on the group distribution pairs (90 & 100, 90 & 110, 100 & 110) using the cSP durations demeaned to their respective subject mean (*Duration* – *Subject Mean*).

EMG spike rate on vs. off was analyzed using a paired t-test, with the rates ‘on’ rate and the ‘off’ rate for each trial paired.

#### Experiment 2, Cortical Excitability

M1_hand_ excitability was analyzed with a paired ranked non-parametric t-test since the distribution was non-gaussian. We performed this using resampling using a script in R (R Core Team, 2021). For null hypothesis testing, permutation was used to create a null distribution of all permutations of before- and after-tUS values swapped within subjects (256 permutations). All 16 medians of each permutation (8 subjects, 2 conditions) were then ranked against one another. The difference of the means of the permuted group ranks for each permutation was used as the values of the null distribution. The value of p equaled the number of permutations in which the absolute difference of the mean ranks was greater than the real absolute difference of the mean ranks.

Confidence intervals were calculated using the bootstrap method (10,000 bootstrap samples). Each bootstrap sample was made by sampling with replacement the 16 real median MEP sizes (8 subjects, 2 conditions). The 95% confidence intervals were set as the 2.5^th^ and 97.5^th^ percentiles of the bootstrap samples’ differences of the group means.

To investigate any association between cortical excitability change and total tUS exposure of M1_hand_, an estimate of total tUS exposure for a participant was calculated with the following formula:

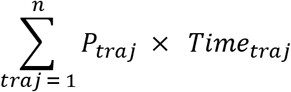

Where ***n*** is the number of tUS trajectories used with that participant, ***P_traj_*** is the pressure (estimated) at M1_hand_ voxel for that trajectory, and ***Time_traj_*** is the time the tUS device was on for that trajectory. Determination of ***P_traj_*** is outlined in k-Wave Simulations. The spearman correlation coefficient (rs) was calculated for these values.

#### cSP Null Distribution

To bootstrap a null distribution of cSP lengths our automated cSP algorithm would find if applied to null, non-cSP data, we used a sliding window approach on non-cSP data. This data was real EMG traces collected during tonic contraction by the same subjects and sessions as Experiments 1 and 2— specifically, the one-second tonic contraction trials collected during tUS exposure. tUS trials were deemed valid as null EMG traces since we saw no change in EMG traces between tUS on vs. tUS off (Figure 4, Figure 5, Supplemental Figure 2, Supplemental Figure 3, Supplemental Figure 4). For every null trial, the cSP algorithm searched for a silence duration from a window centered from each 0.001-second interval from 0.05 to 0.95 s. The first and last 0.05 s were excluded to avoid edge artifacts. All trials, subjects, and sliding-window increments were grouped into a single distribution (686,457 sliding window samples).

## Results

### Experiment 1

#### TMS cSPs

Single-pulse TMS was performed over left M1_hand_ during tonic contraction of the FDI muscle (n = 10). TMS was delivered at 90%, 100%, and 110% aMT. cSP duration increased with TMS intensity (ANOVA: F_2,16_: 26.31, p < 0.001; Welch’s t-tests: p < 0.001, all pairs) (Figure 3, Supplemental Figure 9). The aMT threshold for one subject (sbj11) was set mistakenly low, resulting in TMS intensities lower than intended and therefore elicited very few cSPs.

**Figure 3.**
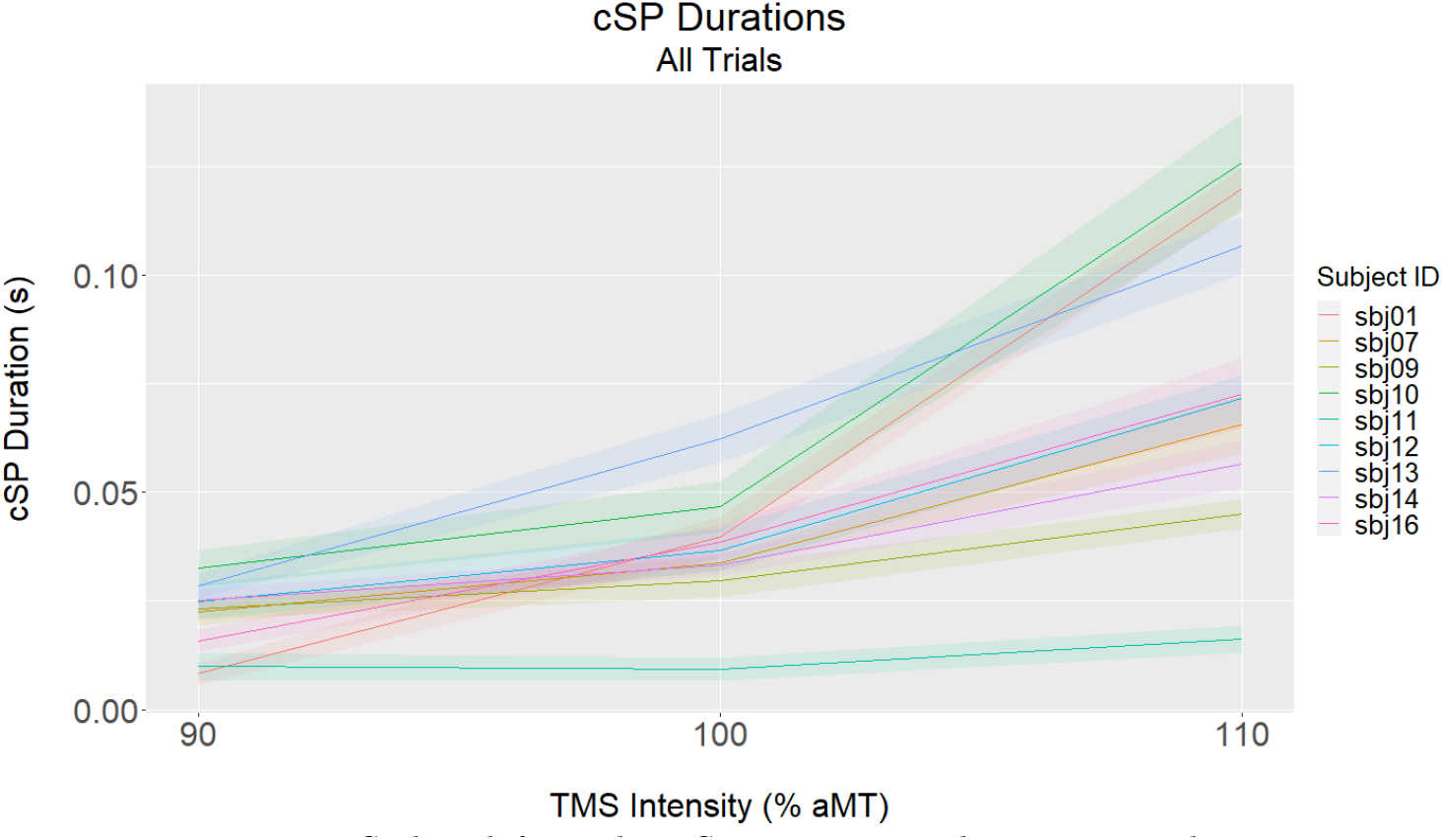
cSP length for each TMS intensity, per subject. Line: median. Ribbon: standard error of the mean. One subject (sbj08) with whom resting motor threshold was used is not shown here (see Supplemental Figure 8).

We also examined the size and presence of MEPs preceding cSPs. For trials with overt MEPs, the lengths of the subsequent silent periods were noticeably longer than would be seen by chance (Supplemental Figure 5). Among trials with a peak within the expected 10-ms time window, trials with a peak smaller than the standard MEP peak-to-peak amplitude threshold of 0.5-mV, henceforth referred to as “stub” trials (Supplemental Figure 10), mostly showed silences within lengths that would be seen by chance (Supplemental Figure 5). However, some “stub” trials did show long silence durations on par with those of overt MEP cSPs. Lastly, trials in which there was *no* peak within the 10-ms time window showed silence durations within what would be seen by chance, with only one of these trials showing a silence above the 95^th^ percentile of the null distribution.

#### tUS

Single-burst tUS was performed over left M1_hand_ during tonic contraction of the FDI muscle (n = 10). The 300-ms or 500-ms bursts were delivered at three trajectories per subject, one of which was also the trajectory for TMS.

No overt silent periods were visible during single trials of tUS stimulation. To investigate whether tUS caused any suppression of the EMG trace, we investigated the height of the EMG traces (area-under-the-curve, AUC) (Supplemental Figure 2), the lengths of the intermittent contraction silences, the rate of EMG peaks (Supplemental Figure 3), and the spectral components of the EMG signals. While a drop in signal power of the spectral components occurs due to TMS-induced cSP, no spectral changes are visible during tUS trials (Figure 4). The same disparity is seen comparing the length of silences in the EMG signal, with a clear rise in mean silence period in response to TMS-induced cSP but no change in response to tUS (Figure 5), For comparisons performed within tUS trials, the height of the EMG traces showed no difference directly before versus after tUS onset (150-ms epochs before vs. after tUS) (Supplemental Figure 2). The rate of EMG peaks while tUS was on vs. off also showed no significant difference (Supplemental Figure 3, Supplemental Figure 4), with a paired t-test confirming there was only a small but statistically insignificant tUS effect on rate of EMG peaks (Delta: −0.91 Hz; 95% CI: −1.99, 0.16 Hz; p = 0.095) (Supplemental Figure 7).

**Figure 4.**
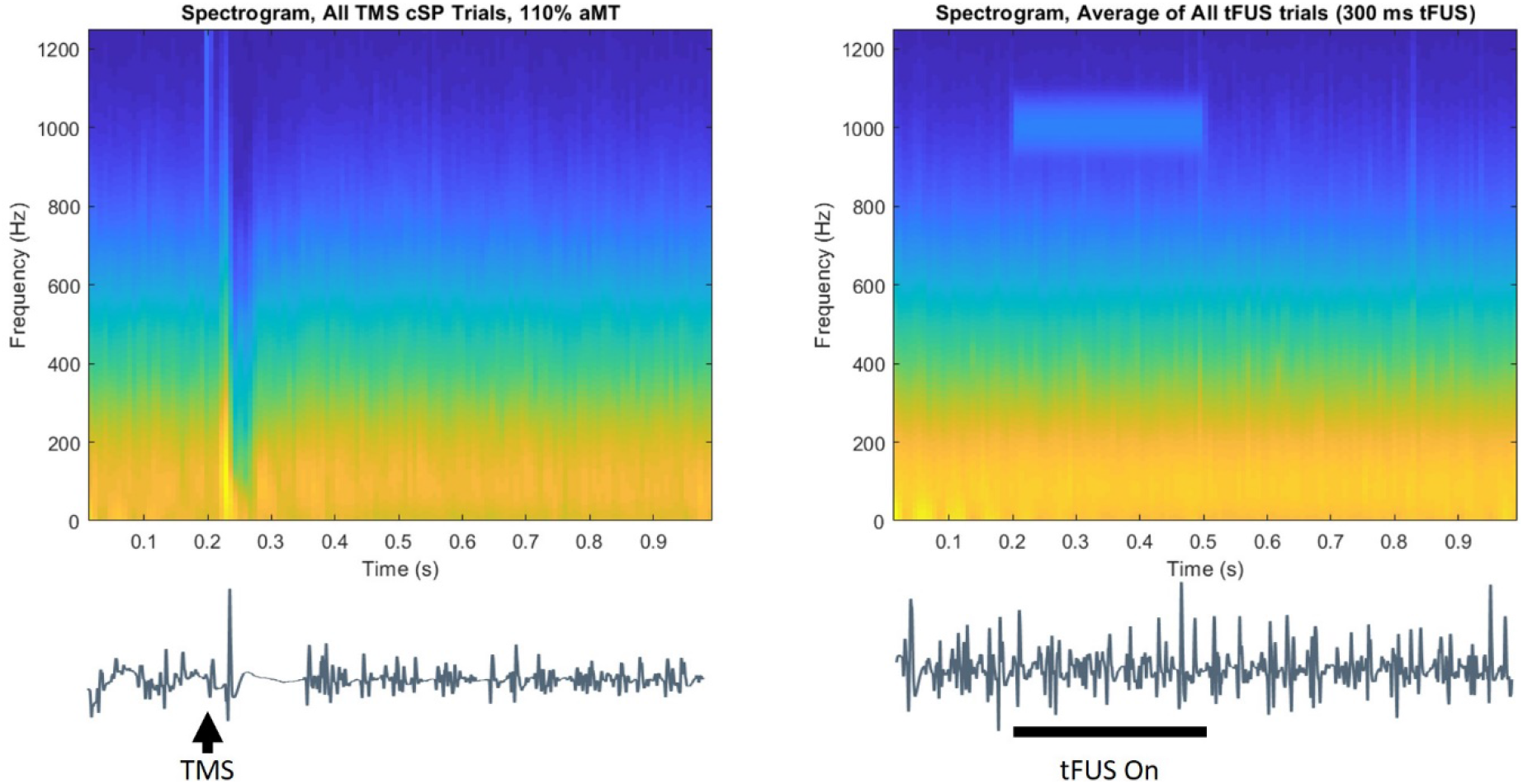
Average spectrograms during tonic contraction. Left) TMS trials. Right) tUS trials (300-ms tUS duration trials only). Signal around the 1000-Hz range from 0.2-0.5 s during tUS trials is noise recorded from the amplifier. This frequency component matches the pulse repetition frequency. Example EMG traces placed below the spectrograms illustrate timing.

**Figure 5.**
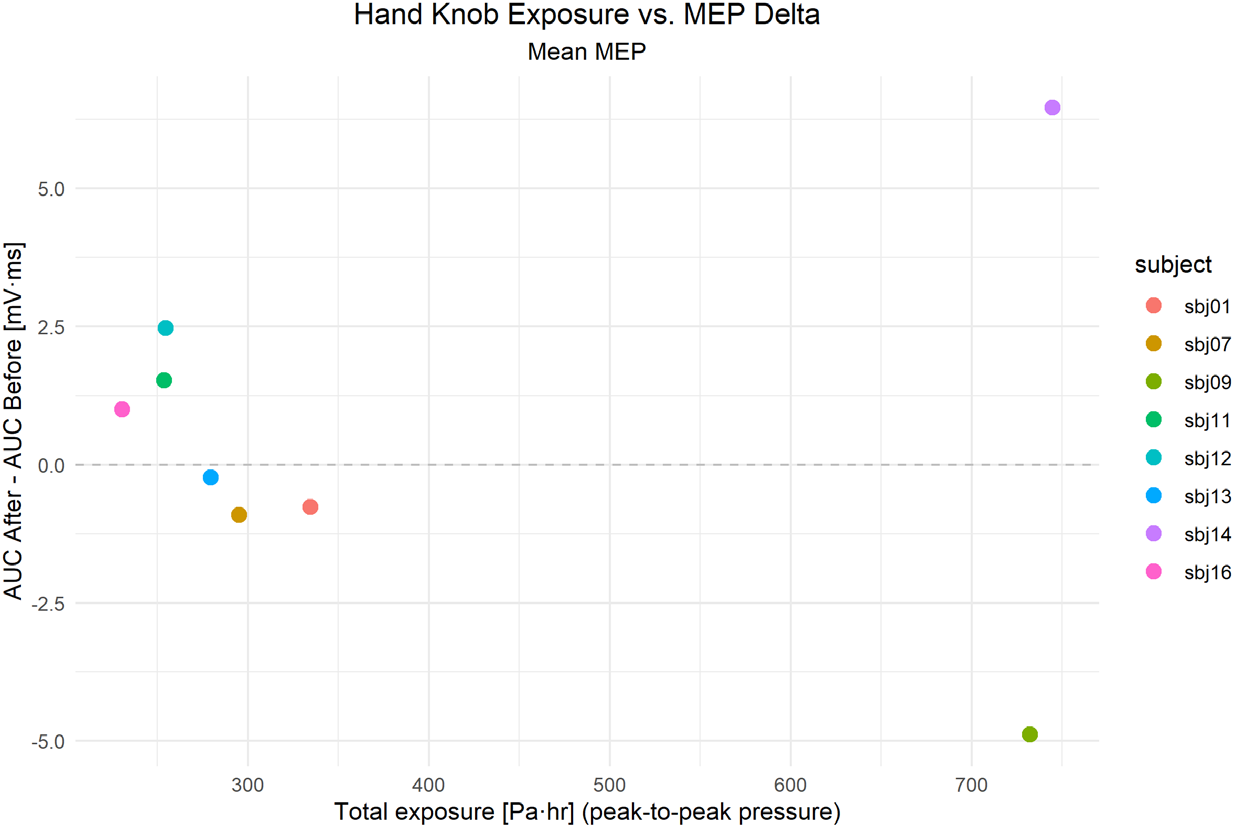
Estimate of total exposure of M1_hand_ to tUS cumulatively during the session (horizontal axis) compared to the change in cortical excitability, as measured by TMS-evoked MEP (vertical axis). r_s_ = −0.21.

### Experiment 2

#### Cortical Excitability

Cortical excitability was gauged before and after exposure to tUS by recording MEPs from single-pulse TMS over M1_hand_ (n = 8). Both the pre-tUS and post-tUS measurements (1-min post-tUS) consisted of 20 suprathreshold TMS trials with an intertrial interval of 10 ± 2 s. The size of TMS-induced MEPs did not vary between before- and after-tUS conditions, according to a ranked paired non-parametric t-test (Figure 8) (Delta: −0.64 mV-ms; 95% CI: −2.39, 0.84 mV-ms; p = 0.51).

#### Exposure vs. Excitability

To investigate the variability that was present among cortical excitability responses, we compared subjects’ cortical excitability change to total estimated tUS exposure in the session. tUS exposure estimates were made using acoustic simulations in models that matched each experimental setup, with skull data computed from structural MRI of the tUS participants. These data showed no obvious correlation between M1_hand_ exposure and cortical excitability change (n = 8), with a spearman correlation coefficient of −0.21 (Figure 6).

**Figure 6.**
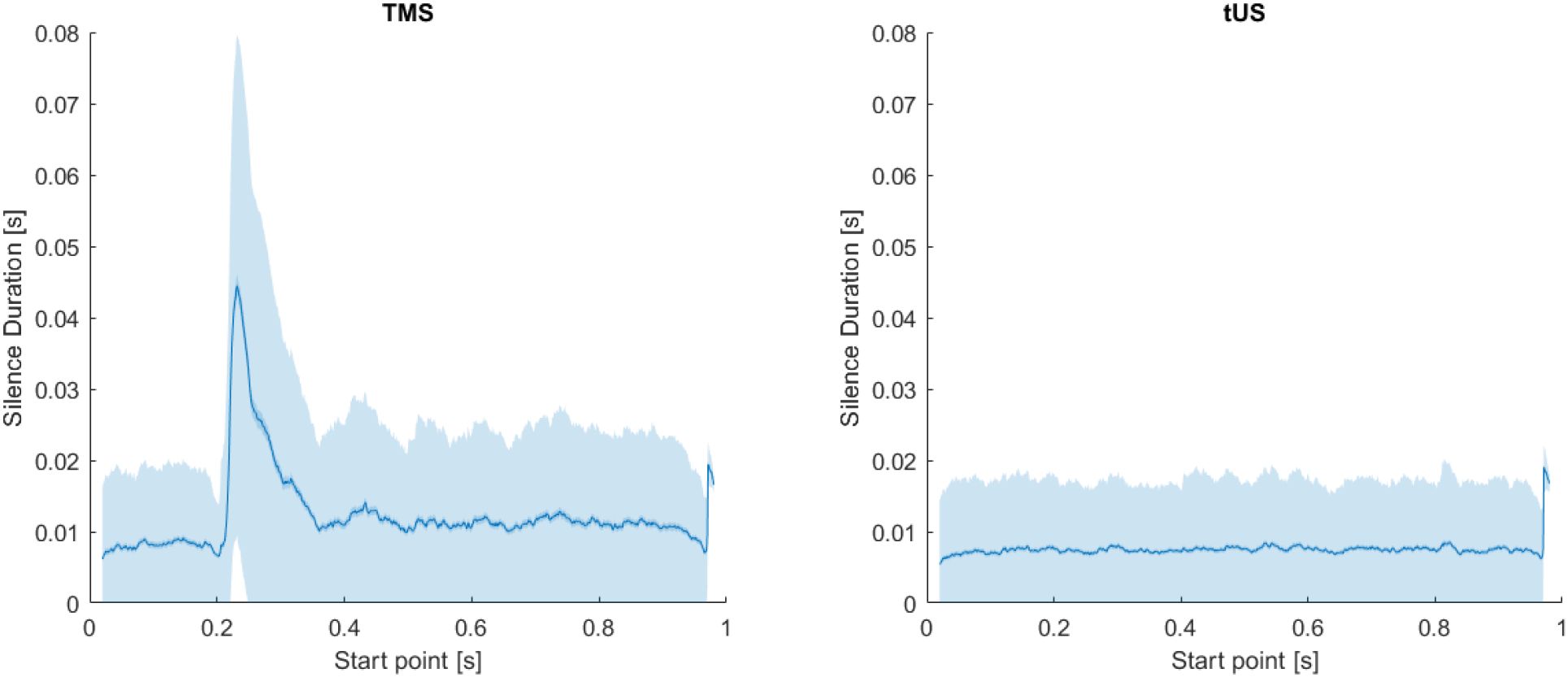
Silence durations across trials, using a sliding window. Duration is as measured from the start point of a given sliding window iteration. Left) TMS; Right) tUS. X-axis: Timepoint measured from. Middle Trace: Mean silence duration. Inner Margin: SEM. Outer Margin: STD

### Acoustic Simulation

Acoustic simulation results suggest we very accurately ‘hit’ targets we were aiming at (Figure 7). tUS produced pressures in an ellipsoid focus, with a mean FWHM with of 4.5 mm (Supplemental Figure 13).

**Figure 7.**
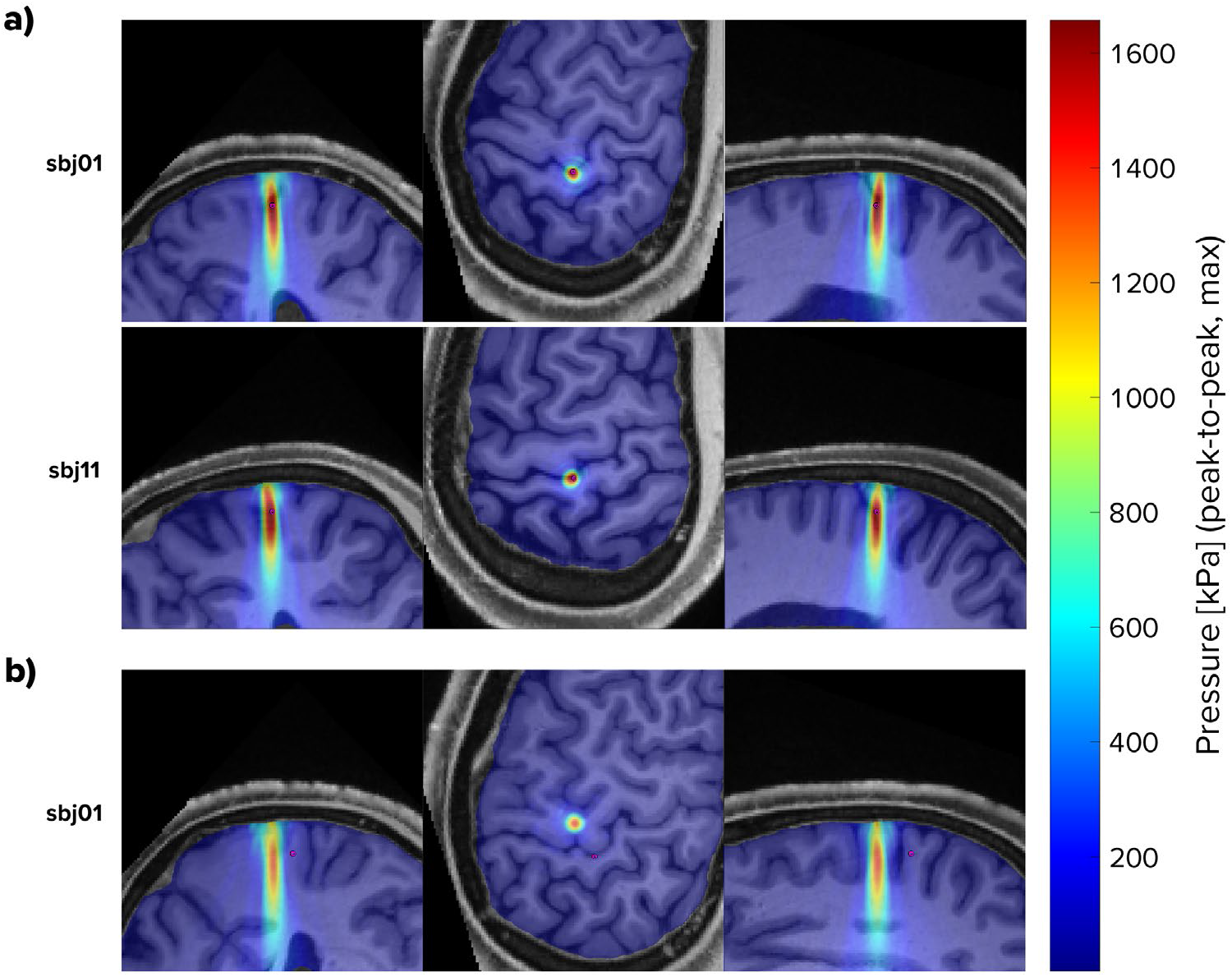
Simulated pressures (examples). a) Two example trajectories that were aimed at the respective subject’s M1_hand_. b) One example trajectory that was aimed at the respective subject’s M1_hand_.

**Figure 8.**
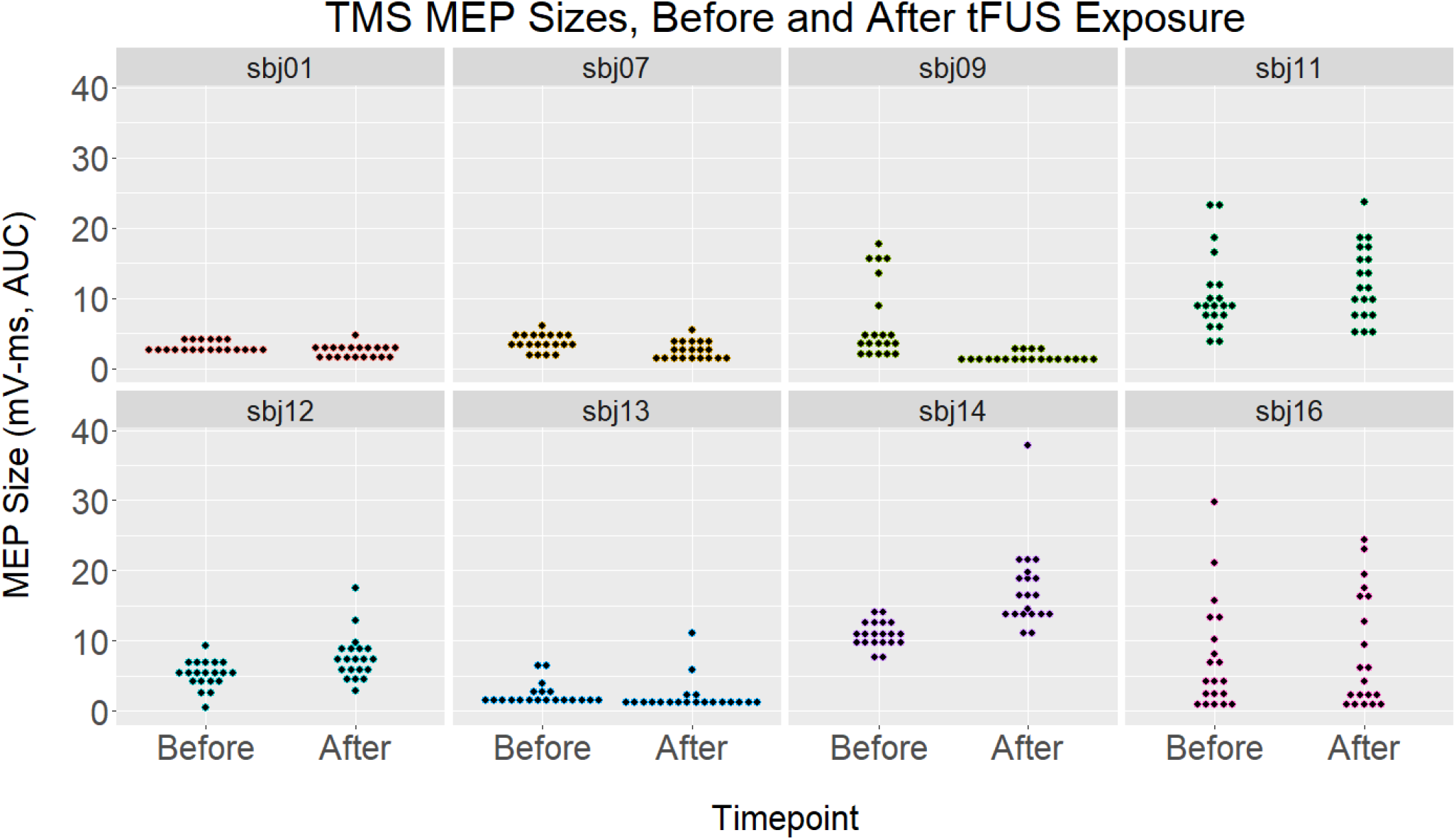
TMS MEP sizes, before and after tUS. Each subplot contains data from one subject: trials before and after tUS on the left and right, respectively. Each point marks the size of one MEP (area under the curve, mV-ms). Points are organized into vertical-axis bins to aid in visualization.

## Discussion

### Experiment 1

#### No tUS-MEPs

We were unable to elicit tUS-induced MEPs at safe intensities. This was the expected outcome, given the lack of MEPs in previous human tUS studies (Ai, Bansal, Mueller, & Legon, 2018). Recent animal model work suggests that motor activation via ultrasound stimulation of motor cortex may not be feasible, proposing that previously reported motor contractions in anesthetized animals likely relied on auditory mechanisms (Guo et al., 2018; Sato, Shapiro, & Tsao, 2018). We did not investigate for potential tUS-MEPs during rest beyond a single pilot subject (data not shown). No tUS-induced MEPs appeared during active contraction trials either.

#### Cortical Silent Period, TMS

Our data found no cSPs that occurred without a preceding TMS-evoked MEP (Figure 6), despite structuring the study to facilitate a high prevalence of near-threshold MEPs. As such, this contradicts claims in the literature that TMS-induced cSPs can occur without an MEP (Classen & Benecke, 1995; Davey et al., 1994; Hupfeld, Swanson, Fling, & Seidler, 2020). One explanation for this difference is that a small “stub” MEP could precede reported “MEP-less” cSPs, with its amplitude not surpassing the amplitude of tonic contraction. This is supported in these data by the consistent appearance of an EMG peak within the latency window expected for TMS-evoked MEPs.

If this MEP-cSP dependency is true, this could suggest that cortical silent period is dependent on the recruitment of M1 motor units. TMS preferentially depolarizes axons (Lazzaro, Ziemann, & Lemon, 2008; Lefaucheur et al., 2014; McIntyre & Grill, 2002; Nowak & Bullier, 1998), while tUS has been proposed to preferentially affect inhibitory neurons (Kim et al., 2014; Legon et al., 2014; Nguyen et al., 2020; Plaksin et al., 2016; Rinaldi et al., 1991). If these assumptions are true, this could explain why tUS struggled to silence corticospinal output.

To be clear regarding the notion of “stub” trials within our TMS data, we do not believe all TMS trials classified as a “stub” by the algorithm are MEPs. Rather, we believe there are two underlying distributions that fall under the “stub” designation. The first: trials in which there is a TMS-evoked MEP that is shorter than the standard threshold (0.5 mV). The second: trials in which there was an EMG peak produced *by chance*—created by a peak in tonic muscle EMG activity that fell within the expected time window (Supplemental Figure 10).

We must also note: For cSP length determination, we took a conservative approach on brief EMG activity flanked by periods of silence, referred to in the literature as late excitatory potentials (LEPs) (Butler, Petersen, Herbert, Gandevia, & Taylor, 2012; Kallioniemi et al., 2015; Wilson, Thickbroom, & Mastaglia, 1995). Of the two silent periods flanking an LEP, we included only the first silent period when measuring cSP duration. Since these LEPs appear heuristically as short EMG disruptions of a longer cSP, a visual inspection of the data suggests that ignoring these LEPs would have resulted in less variable cSP durations within blocks (Supplemental Figure 6). For comparison, these LEPs have at times been ignored in past by-hand cSP measurements (Hupfeld et al., 2020).

#### Cortical Silent Period, tUS

Single-burst tUS of the hand area of left motor cortex did not affect tonic muscle contraction of the FDI muscle. Specifically, there were no deviations in gap duration between tonic muscle spikes (Figure 5), spectral components (Figure 4), or prevalence of EMG peaks (Supplemental Figure 3, Supplemental Figure 4). This is in sharp contrast to the lengthy silent periods from single-pulse TMS.

Since these data revealed no time-locked tUS effects, this leaves open the question whether tUS affects cortical motor circuitry *at these parameters*. Looking to the TMS literature for insight, we know that it *is* possible to induce detectable excitation of motor cortex without a measurable peripheral effect. Specifically, electroencephalography (EEG) recordings during subthreshold single-pulse TMS show significant TMS-evoked potentials, despite a lack of a peripheral MEP (Gordon, Desideri, Belardinelli, Zrenner, & Ziemann, 2018). Given this, it is possible there were tUS effects that did not interact with corticospinal projections—making them undetectable by EMG and therefore undetectable by our experiment. While EEG is not necessarily a more sensitive readout to study all mechanisms, such as for certain corticospinal excitability experiments (Desideri, Zrenner, Gordon, Ziemann, & Belardinelli, 2018), methods like EEG and fMRI that record cortical effects directly could provide a more complete picture of time-locked tUS effects in future investigations.

### Experiment 2

#### Cortical Excitability

Our data showed no group difference in M1 excitability in response to tUS, as indicated by no change in MEPs evoked by TMS at the same M1 trajectory. Our data differ from data by Gibson and colleagues that showed increased excitability of M1 after ultrasound exposure (Gibson et al., 2018). There are study design differences that could have driven this disparity. The first is the tUS device used. While we used a 500-kHz single-element focused transducer, *Gibson et. al 2018* used an imaging ultrasound device, which consisted of an array of 80 transducer elements emitting frequencies in a range between 1.53 and 3.13 MHz. This frequency range is noteworthy because acoustic attenuation increases as a function of acoustic frequency (Hayner & Hynynen, 2001; *The Safe Use of Ultrasound in Medical Diagnosis*, 2012; White, Clement, & Hynynen, 2006). *Gibson et al. 2018* cited papers that used ultrasound imaging devices to image through the skull as evidence of the device’s validity for use over M1. However, all studies they cited placed the device over the temporal window—an area of the skull that is significantly thinner than that over M1 (temporal window: ~3 mm; parietal bone: ~6 mm) (Kwon, Kim, Kang, Bae, & Kwon, 2006; Mahinda & Murty, 2009). In fact, measurement of ultrasound propagation through ex-vivo human parietal skull shows that little to no energy is transmitted at frequencies above ~1.5 MHz (Hynynen & Jolesz, 1998; O’Brien, 2007; Pichardo, Sin, & Hynynen, 2010; White et al., 2006).

The second noticeable difference between the two studies is the stimulation protocol. While *Gibson et al.* delivered constant exposure to an ultrasound imaging protocol for 2 minutes, we delivered separate bursts of ultrasound (duration: 300-500 ms each) with long gaps between bursts (8-12 s inter-burst interval)—an interval that was chosen to support the primary aim of this study: investigation for silent periods. Our ~10-second interval is very slow compared to repetitive TMS protocols used to modulate cortical excitability, and it ventures into the intertrial interval range suggested for use to avoid central habituation effect in sensory stimulation studies.(Baumgärtner, Greffrath, & Treede, 2012; Greffrath, Baumgärtner, & Treede, 2007; Warbrick, Derbyshire, & Bagshaw, 2009) If neuromodulation is the aim, future tUS studies may want to use more compressed protocols with shorter inter-burst intervals. A compressed approach was shown to successfully affect tUS targets in non-human primates, as shown by the reduction of resting-state fMRI connectivity following a 40-s tUS protocol (pulse repetition frequency: 10 Hz; pulse length: 30 ms) (Folloni et al., 2019; Verhagen et al., 2019). These non-human primate studies showed effects lasting up to two hours after stimulation. However, they also used tUS intensities, 24.1-31.7 W/cm^2^, that were significantly higher than the levels used here or any other human tUS study (human max.: 4.9 W/cm^2^ in *Legon et al. 2014*). While histological examination in these studies revealed no microstructure damage, the protocol in question still corresponds to a mechanical index of ~3.6—higher than the 1.9 maximum allowed by the FDA for diagnostic imaging (Şen, Tüfekçioğlu, & Koza, 2015; *The Safe Use of Ultrasound in Medical Diagnosis*, 2012). Given that the most robust tUS effects seem to occur at high intensity levels, future studies will need to carefully explore whether consistent, behaviorally relevant tUS effects are feasible at intensities safe for human exposure. Replication of repetitive tUS protocols, at lower intensities, will likely be the first step.

#### Exposure vs. Excitability

With this small sample size (n = 8), we saw no correlation between M1_hand_ exposure and cortical excitability change, though the calculation of this correlation was also low-powered. This conclusion is to be expected since this study was not designed to investigate such a correlation, with M1_hand_ exposure effectively stratified into two levels depending on whether the M1_hand_ target was used once or twice (i.e. whether it was the primary NIBS target, “TMS target”). Studies that wish to investigate potential correlation effects of exposure levels would need to expose participants at a variety of different levels and have a larger sample size than used here to increase statistical power compared.

While we performed acoustic simulation on 907 cm^3^ volumes (~300-350 cm^3^ of which were grey or white matter), we still chose to tabulate cumulative exposure for a single location: M1_hand_. M1_hand_ was chosen because it is the most reasonable small-volume, easily identifiable cortical area that is known to play a direct role in hand muscle contraction. It is, however, possible that directly targeting M1_hand_ may not be the ideal choice for modulating voluntary muscle contraction with tUS. This notion of placement is especially important given the spatially precise nature of tUS compared to a TMS or transcranial electrical stimulation—especially with a small, focused ultrasound transducer as used here.

### tUS and M1

While there could be multiple causes, one potential explanation could lie in cytoarchitectural differences between brain regions. Crucially, motor cortex has significantly lower neuronal density compared to somatosensory and visual cortex (Atapour et al., 2019; Beaulieu & Colonnier, 1989; Collins, 2011; Collins, Airey, Young, Leitch, & Kaas, 2010). As such, for a tUS pressure field of a given size, the number of individual neurons that fall within the focus would be lower in M1 compared to primary somatosensory (S1) or primary visual cortex (V1). This disparity could leave M1 neurons at a relative disadvantage for reaching thresholds to create detectable systems-level effects from tUS exposure. Additionally, the inherent cytoarchitectural and circuitry differences between ‘output’ cortical regions, like M1, compared to ‘input’ cortical regions, like S1 and V1, could likely have a significant role.

## Conclusion

We performed neuronavigated tUS and TMS of primary motor cortex (M1) in healthy volunteers. We found no concurrent change in finger EMG activity from tUS of M1 during voluntary muscle contraction. We also did not find any consistent effect of tUS M1 exposure on motor cortex excitability, as measured by single-pulse TMS of M1. We performed acoustic simulations using structural MRI of the study participants to estimate the degree and location of ultrasound intracranially. Using these simulations, we were unable to find any correlation between cumulative ultrasound exposure of the M1 hand area and M1 excitability change.

Within the TMS-only data, our data suggest that cortical silent periods (cSP) may be contingent on a motor evoked potential (MEP) occurring at cSP onset, though at times the MEP may elude visual detection due to a small amplitude that does not rise above the level of tonic muscle activity. This finding questions previous reports of cSPs without MEPs (Classen & Benecke, 1995; Davey et al., 1994; Hupfeld et al., 2020).

While the negative tUS results reported here mirror struggles other investigators have shown when attempting to elicit measurable modulation of M1 by tUS, this was also a pilot study with small sample sizes (n = 8; n = 10). As such, clearer results may emerge with larger datasets or changes in methodology.

## Notes

### Competing Interest Statement

The authors have declared no competing interest.

