## Supplemental Figures for "Ultrasound stimulation of the motor cortex during tonic muscle contraction"

Supplement for preprint deposited into *bioRxiv*. This version has not been peer reviewed.
bioRxiv version 1

**Title:** *Ultrasound stimulation of the motor cortex during tonic muscle contraction*
**Authors:** Ian S. Heimbuch, Tiffany Fan, Allan Wu, Guido C. Faas, Andrew C. Charles, Marco Iacoboni
**Institution:** University of California, Los Angeles

### SUPPLEMENTAL FIGURES

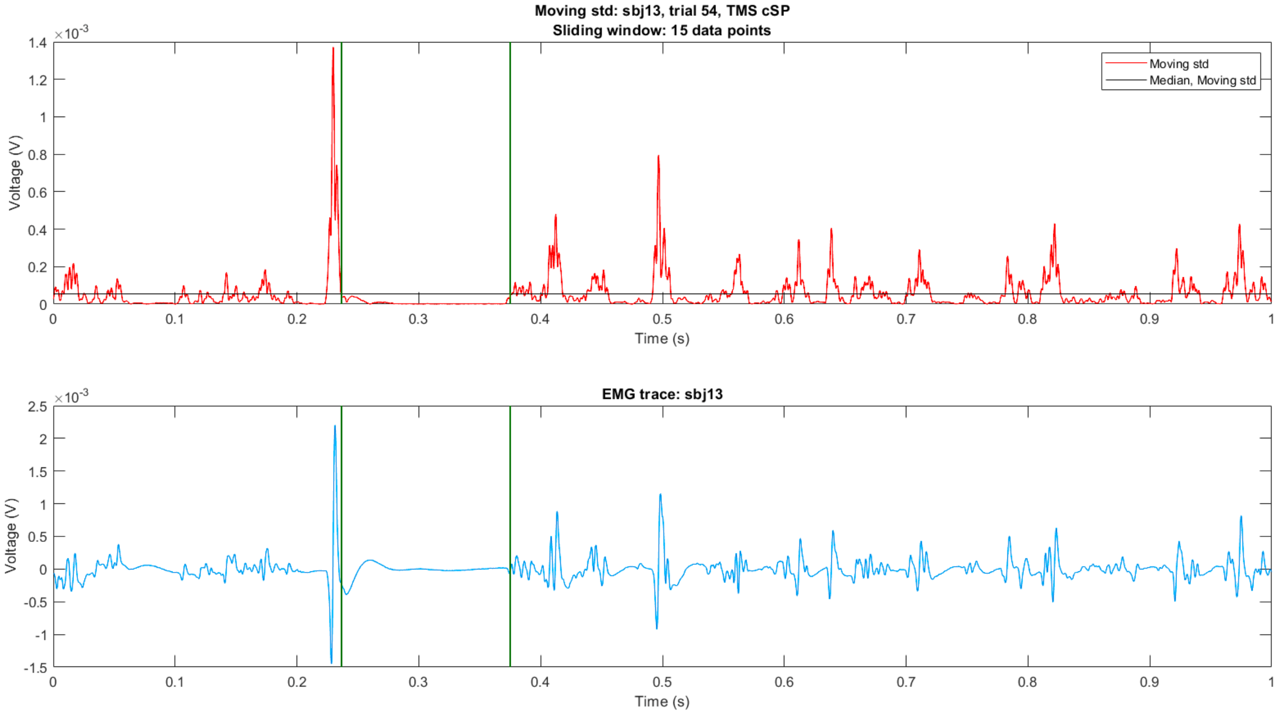


*Supplemental Figure 1. Visualization of the automated cSP detection method. Top) Sliding window standard deviation trace of a single trial EMG trace. The black horizontal line marks the detection threshold. The vertical green lines mark the beginning and end of the detected cSP. Bottom) The original high-pass filtered EMG trace.*

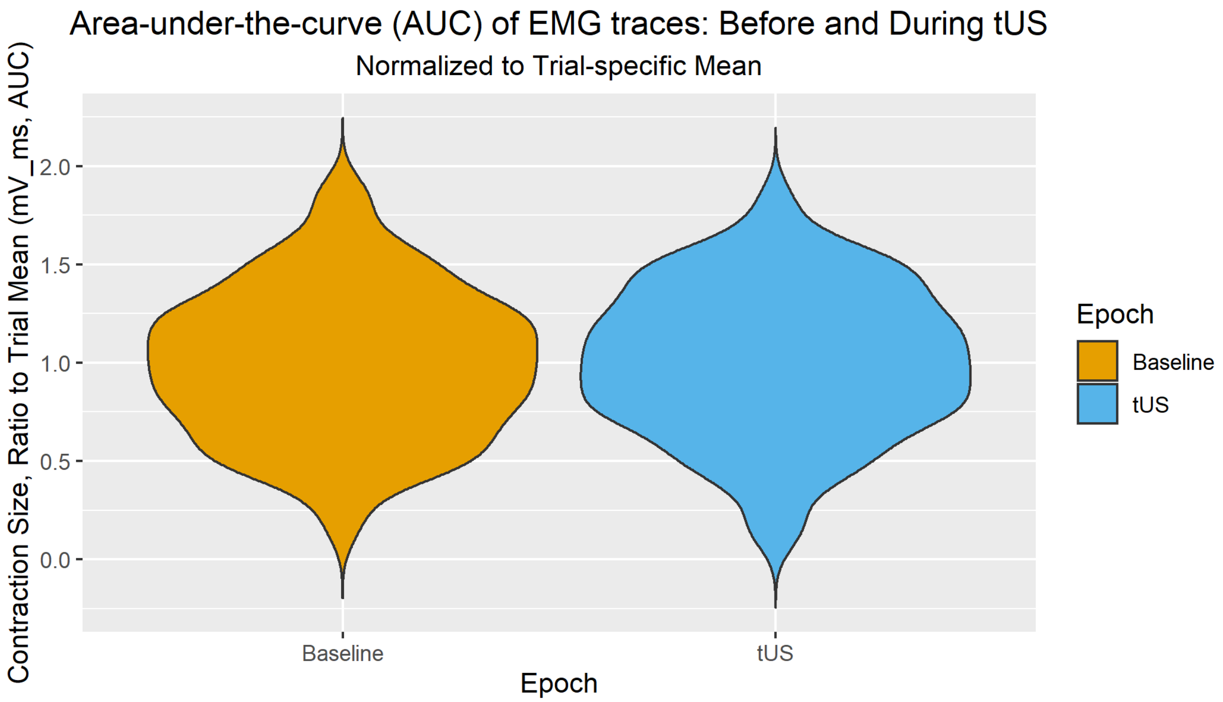


*Supplemental Figure 2. Area-under-the-curve of tUS traces. Level of EMG activity during different sections of tUS trials. Left) Baseline, -200 to -50 ms before onset. Right) 0 to 150 ms after tUS onset (first 150 ms of tUS exposure). Values were normalized via dividing by the trial mean.*



Supplement for preprint deposited into *bioRxiv*. This version has not been peer reviewed.
*Ultrasound stimulation of the motor cortex during tonic muscle contraction.* (bioRxiv version 1)

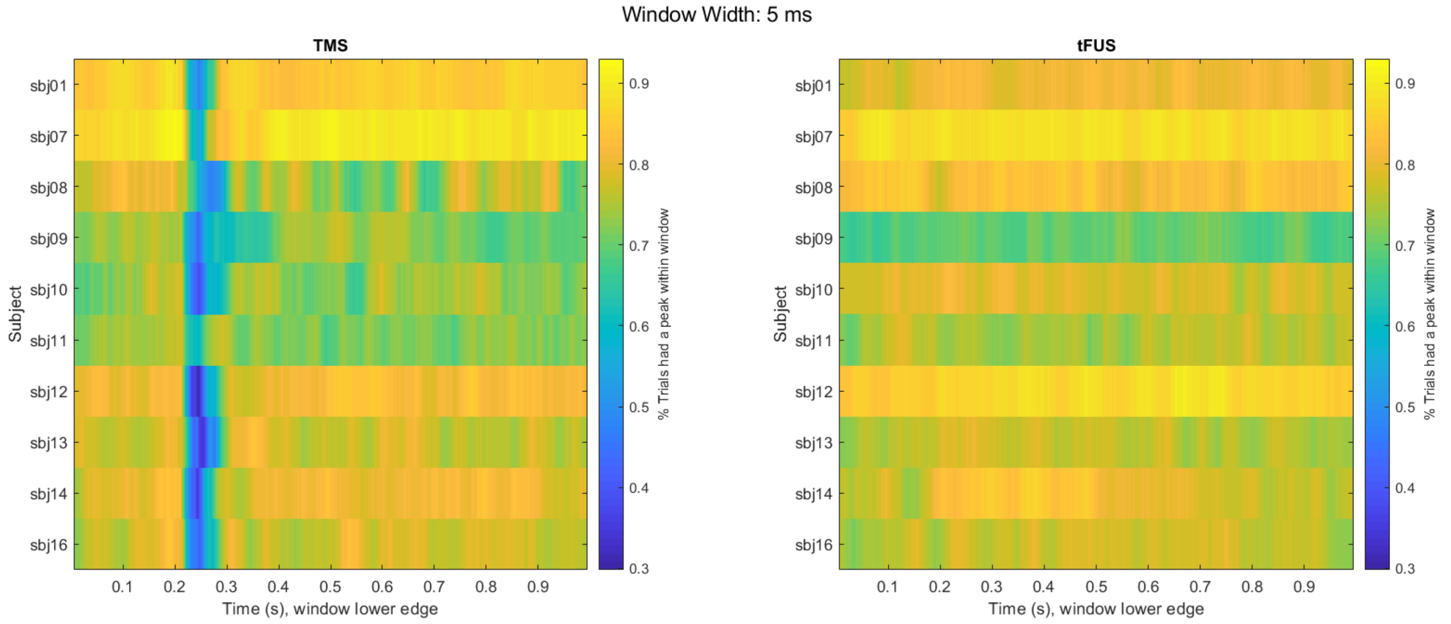


*Supplemental Figure 3. Prevalence of EMG peaks. A sliding window approach (1-ms steps) checked if an EMG peak occurred during a 5-ms time window following that point. Peaks were detected using findpeaks() function in MATLAB. Each row contains data for all trials per subject. Color shows percentage of trials that had a peak during that time window. Percentage data was smoothed with a moving mean (~6-ms window). Left) TMS trials. Right) tUS trials. EMG traces were high pass filtered at 10 Hz.*

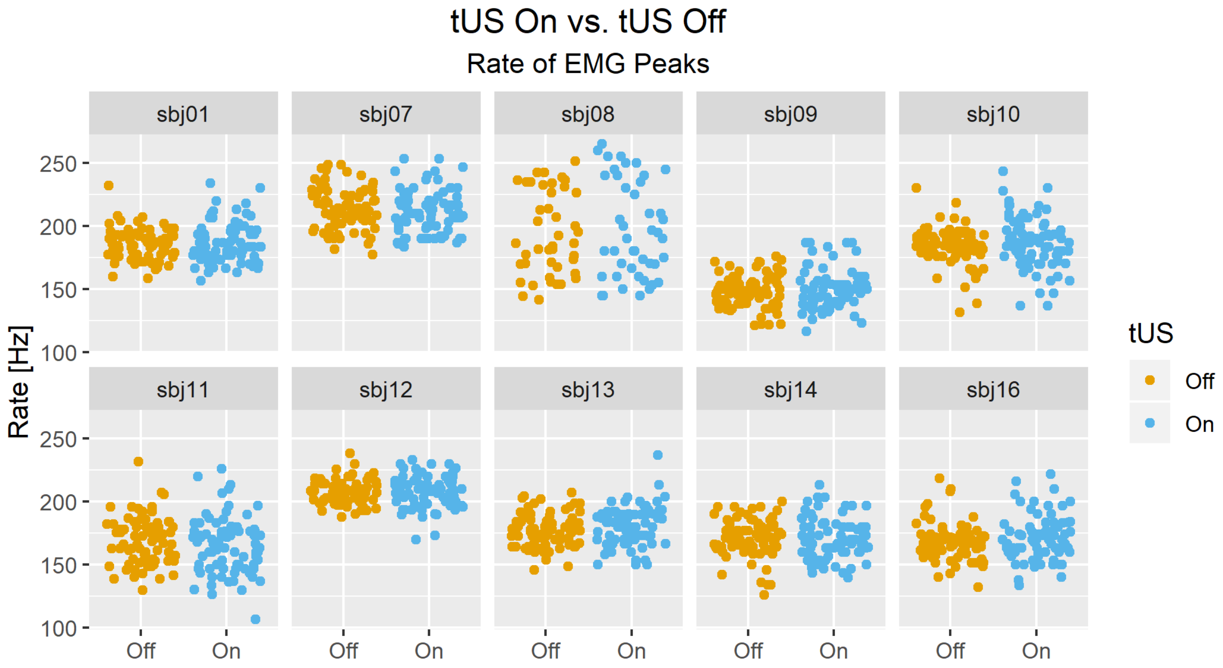


*Supplemental Figure 4. Rate of EMG peaks during a single tUS trial. One dot per trial per condition ('Off' and 'On'). EMG traces were bandpass filtered to 10-800 Hz.*


Supplement for preprint deposited into *bioRxiv*. This version has not been peer reviewed.
*Ultrasound stimulation of the motor cortex during tonic muscle contraction.* (bioRxiv version 1)

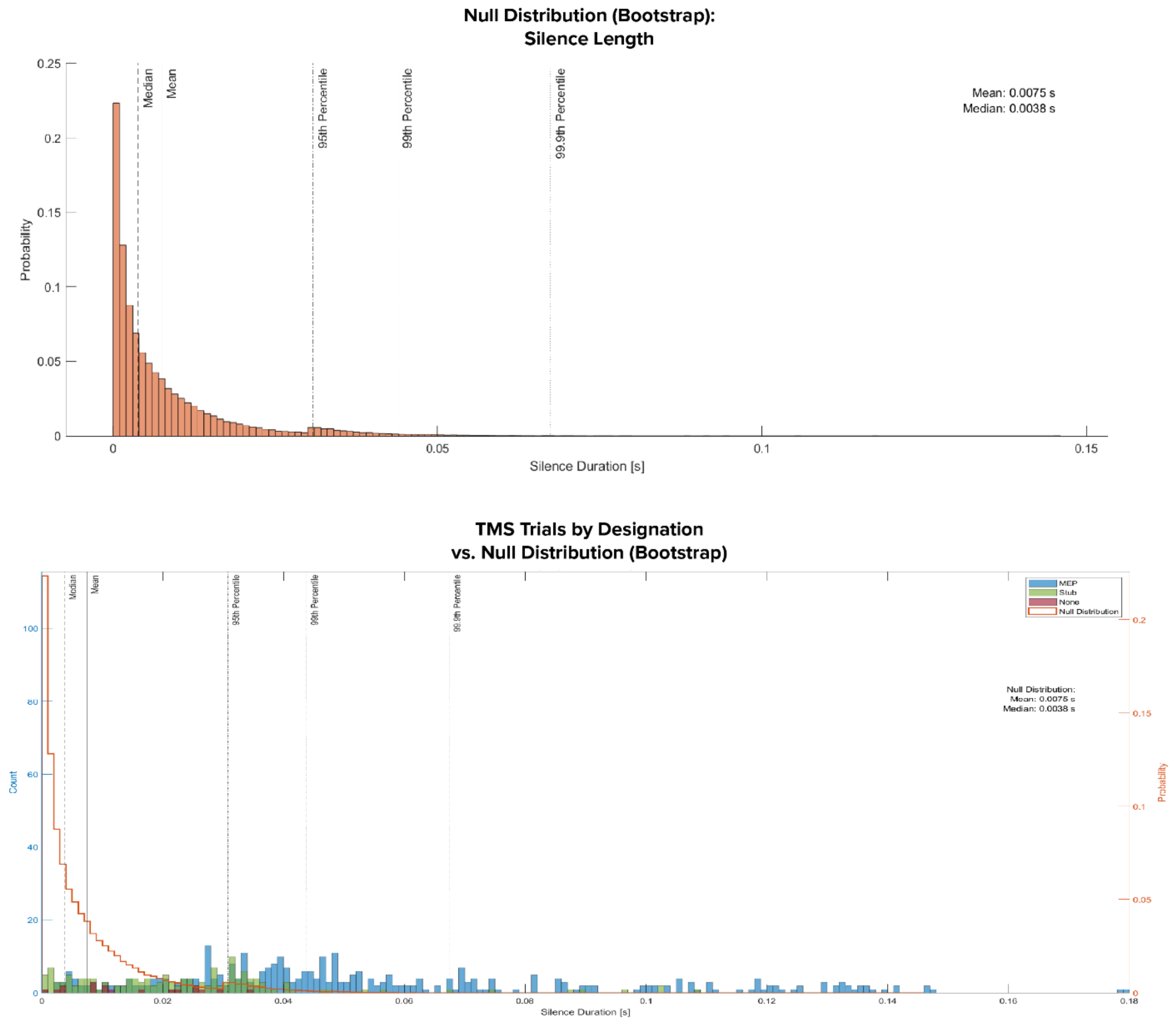


*Supplemental Figure 5. **Top:** Distribution of silence lengths when cSP length algorithm is run from different time points along each EMG from a contracting finger. Sliding window approach was used to bootstrap these values, with a sliding window step size of 0.001 seconds. Histogram bin width 0.001 seconds. "Null" data were the one-second tonic contraction trials during tUS exposure. tUS trials were deemed valid as null EMG traces since we saw no change in EMG traces between tUS on vs. tUS off (Figure 4, Figure 5, Supplemental Figure 4). The first and last 50 ms were removed to avoid boundary effects. 686,457 sliding window samples. **Bottom:** Lengths of silent periods for trials grouped into three categories: a clear MEP was present ("MEP", blue), a small peak that may have been an MEP was present ("Stub", green), and no detectable peak was present ("None", red). Null distribution (see Top) overlayed in orange. MEP: 361. Stub: 151. None: 18.*



Supplement for preprint deposited into *bioRxiv*. This version has not been peer reviewed.
*Ultrasound stimulation of the motor cortex during tonic muscle contraction.* (bioRxiv version 1)

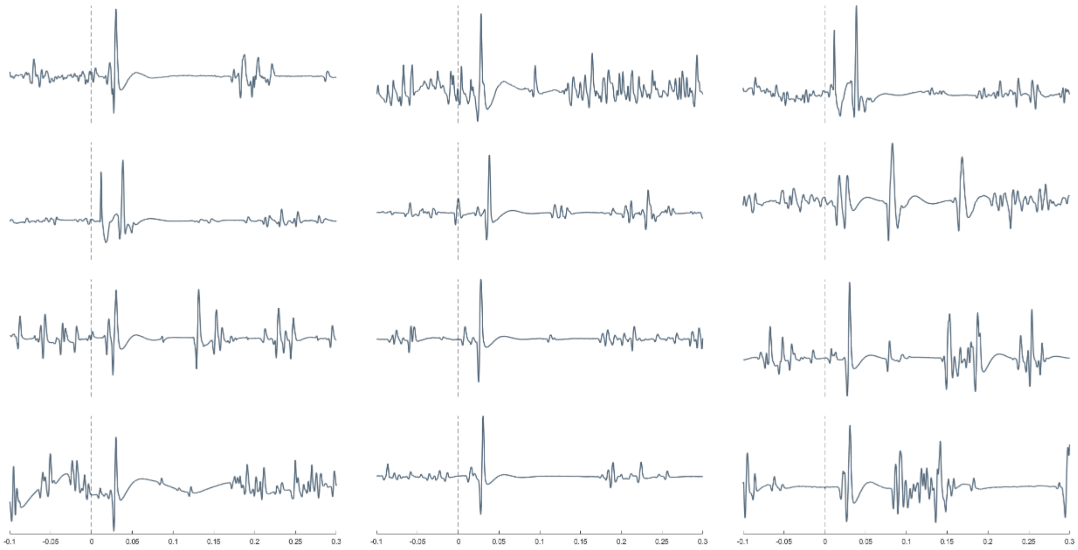


*Supplemental Figure 6. Examples of cSPs with late excitatory potentials (LEPs). X-axis: Time [s]. TMS onset at 0 s. Examples are from multiple subjects.*


Supplement for preprint deposited into *bioRxiv*. This version has not been peer reviewed.
*Ultrasound stimulation of the motor cortex during tonic muscle contraction.* (bioRxiv version 1)

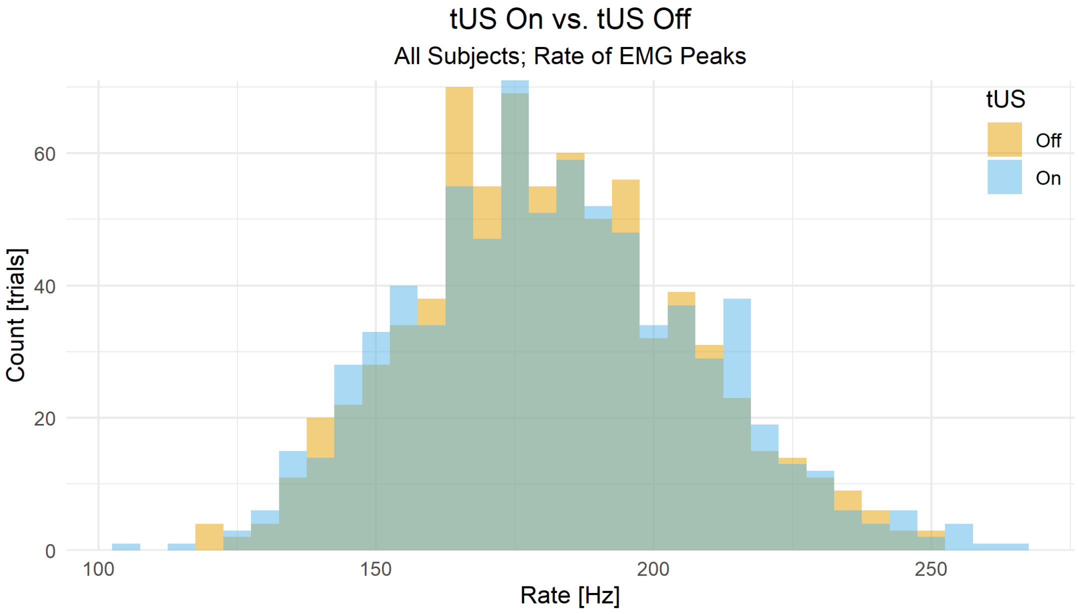


*Supplemental Figure 7. Distributions of rate of EMG peaks during a single tUS trial. All subjects; all trials. EMG traces were bandpass filtered to 10-800 Hz. Same data as shown per subject in Supplemental Figure 4. Difference of the mean rates of EMG peaks were only marginally lower for tUS 'On' vs. tUS 'Off' (Delta: -0.91 Hz; 95% CI: -1.99, 0.16 Hz; p = 0.095; paired t-test).*



Supplement for preprint deposited into *bioRxiv*. This version has not been peer reviewed.
*Ultrasound stimulation of the motor cortex during tonic muscle contraction.* (bioRxiv version 1)

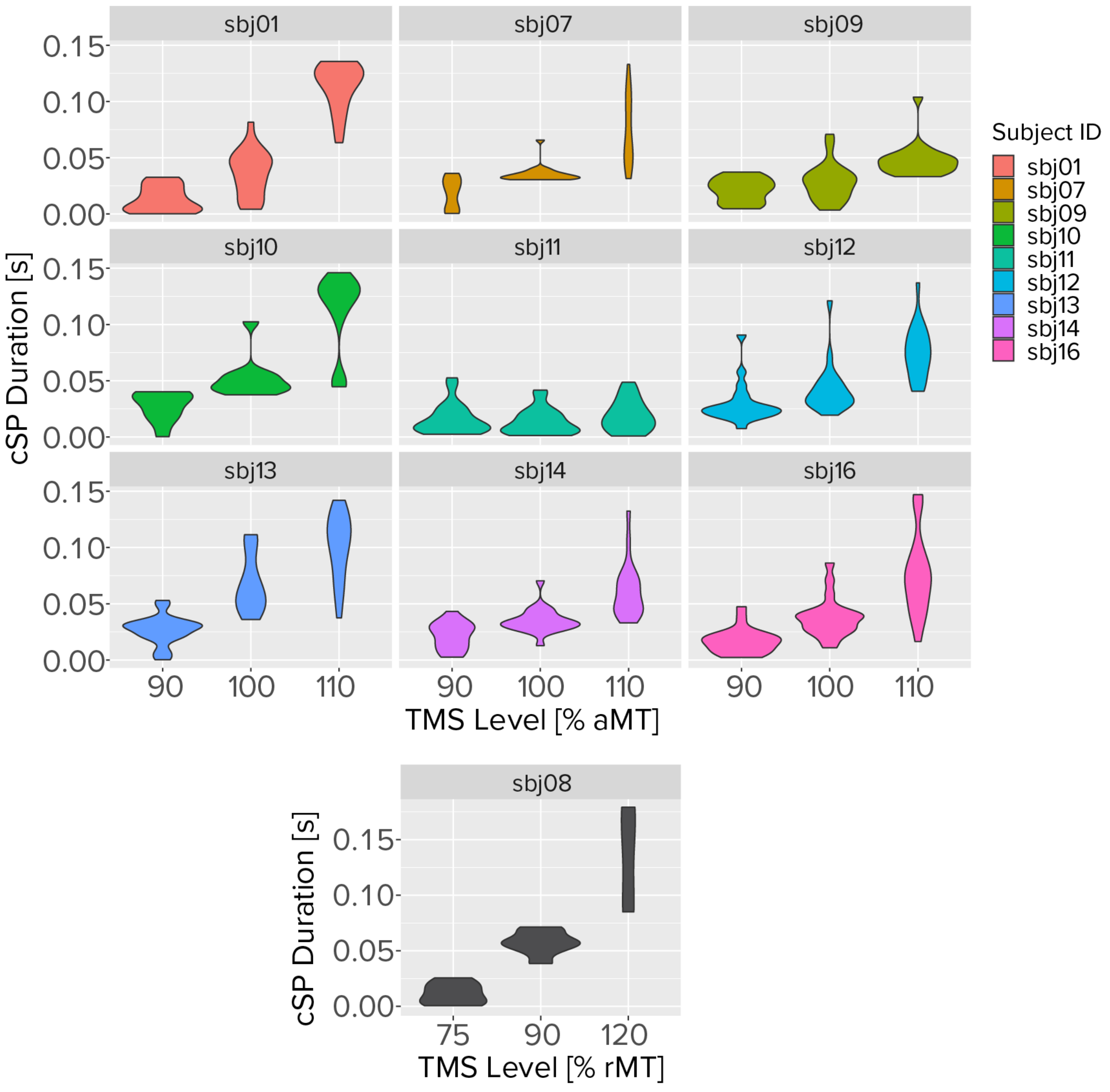


*Supplemental Figure 8. Violin plots of TMS-evoked cSP durations separated by research participant ("Subject ID"). Same data as shown in Figure 3. sbj08 subject was excluded from Figure 3 since their cSP trials were not used in analysis (due to use of different TMS levels). Non-zero silence durations were recorded for 90% aMT for two reasons. First, there are inherent gaps between EMG peaks during tonic contraction, which average ~7.5 ms with our algorithm and our data (Supplemental Figure 5, Top). Second, TMS of M1 results in a distribution of responses, with some trials reaching MEP and cSP threshold while other trials do not (i.e. motor thresholds are never hard cutoffs).*



Supplement for preprint deposited into *bioRxiv*. This version has not been peer reviewed.
*Ultrasound stimulation of the motor cortex during tonic muscle contraction.* (bioRxiv version 1)

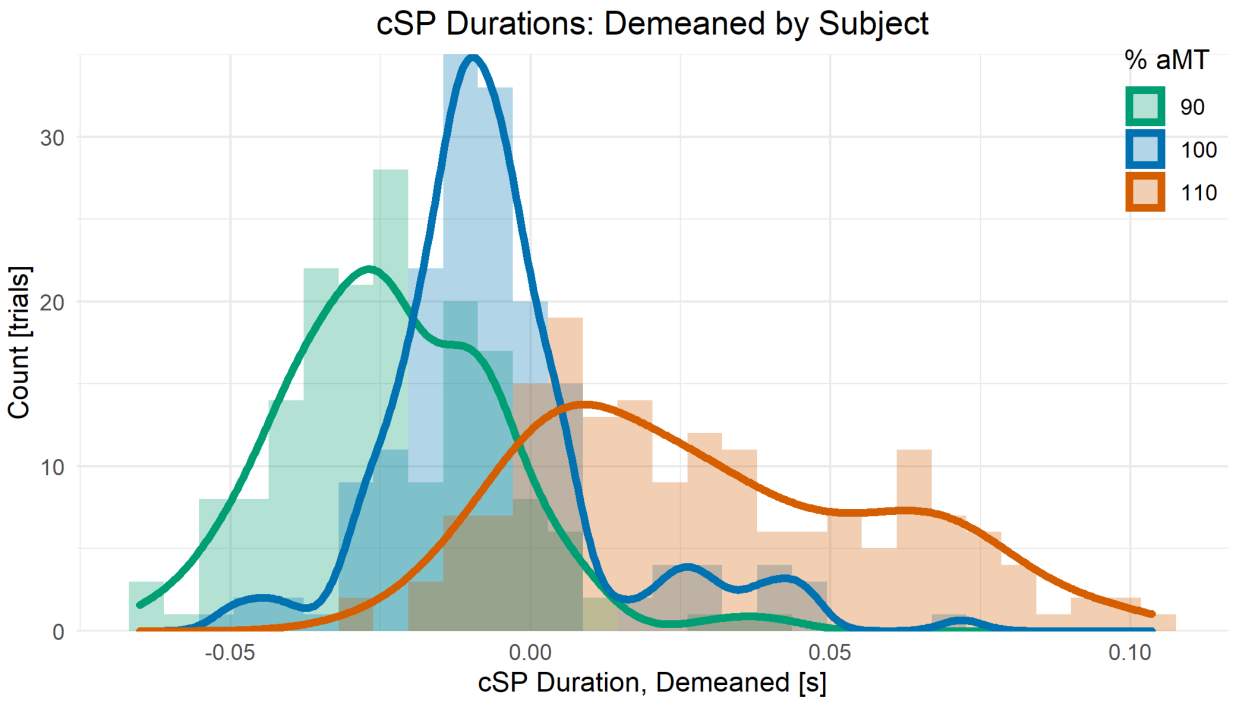


*Supplemental Figure 9. cSP durations demeaned by subject mean. Histograms and density plots shown by % aMT. Welch's t-tests performed as post-hoc tests confirmed cSP duration increased by % aMT (p < 0.001, all pairs). For non-demeaned data see Figure 3 and Supplemental Figure 8. One subject (sbj08) with whom resting motor threshold was used is not shown here (see Supplemental Figure 8).*


Supplement for preprint deposited into *bioRxiv*. This version has not been peer reviewed.
*Ultrasound stimulation of the motor cortex during tonic muscle contraction.* (bioRxiv version 1)

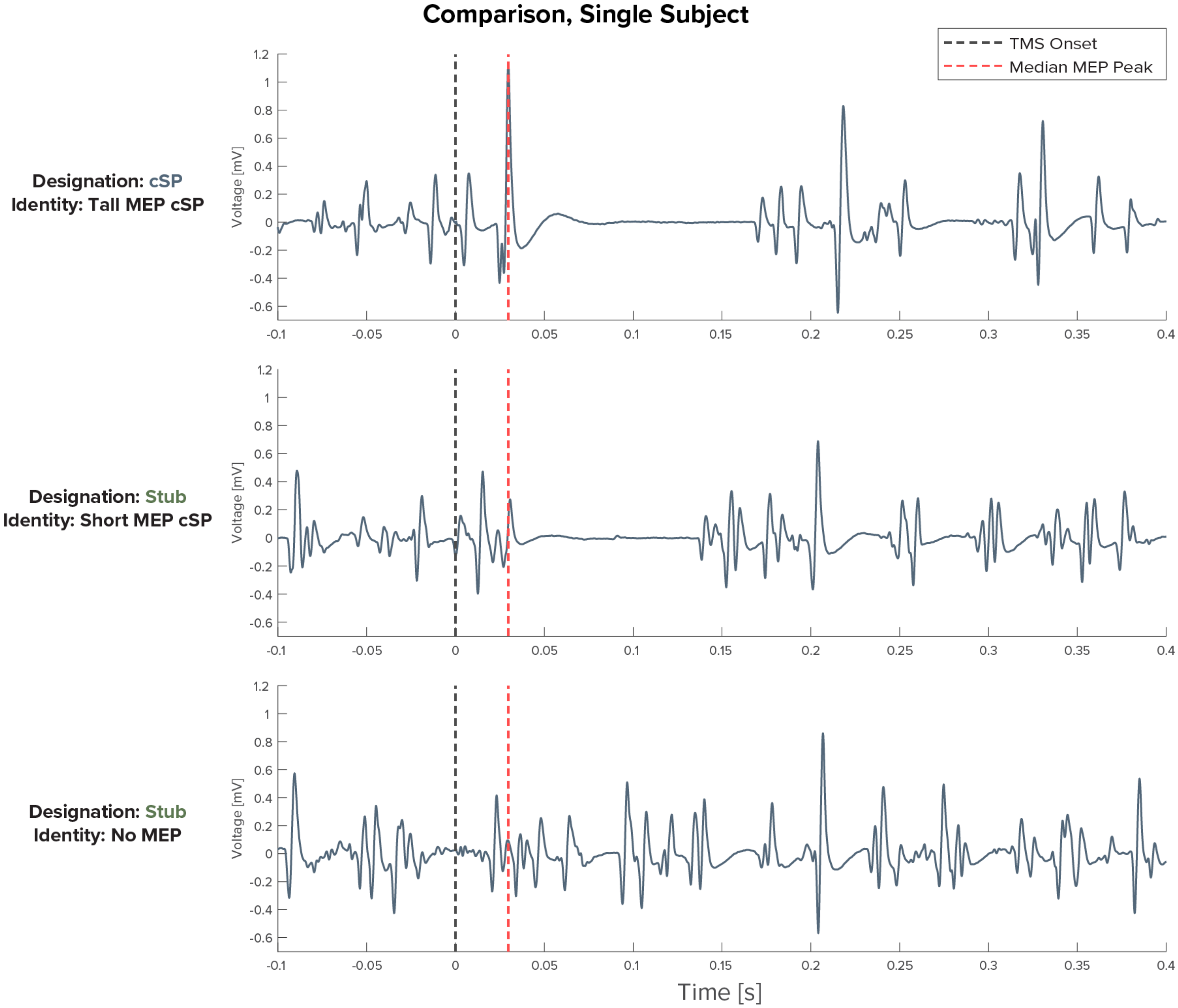


*Supplemental Figure 10. Comparison of three different results from single-pulse TMS during tonic contraction. All three trials are from the same subject. The automated trial designation classified each trial (**Top**, **Middle**, **Bottom**) as: "cSP", "Stub", "Stub". These two distinct examples of scenarios that fall under the "Stub" designation, as determined by the algorithm. This illustrates that there are likely two main distributions of trials that fall under the "Stub" designation. The first: trials in which there is a TMS-evoked MEP that is shorter than the standard threshold (0.5 mV). The second: trials in which there was by chance an EMG peak produced by tonic muscle contraction that fell within the expected time window.*



Supplement for preprint deposited into *bioRxiv*. This version has not been peer reviewed.
*Ultrasound stimulation of the motor cortex during tonic muscle contraction.* (bioRxiv version 1)

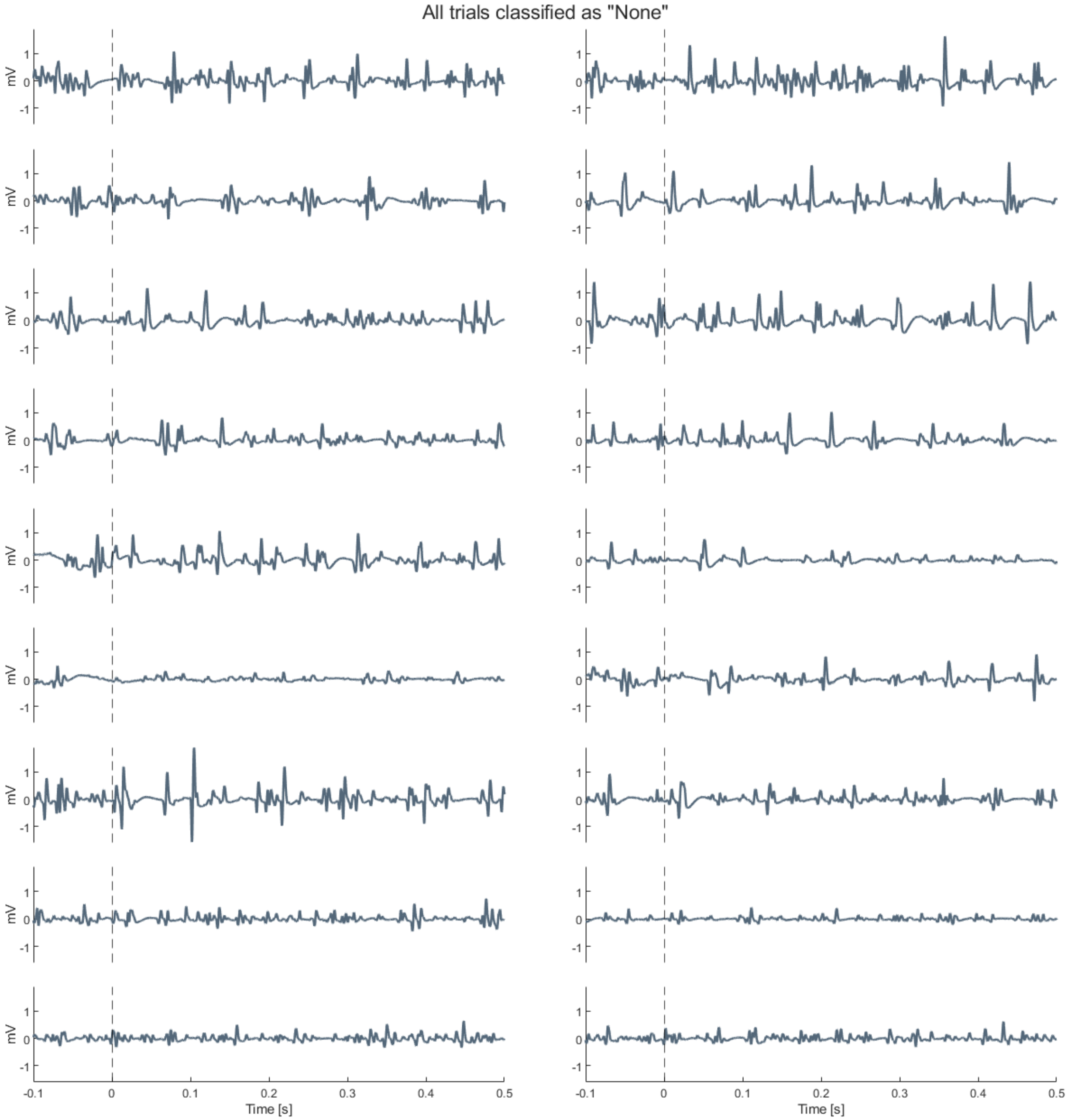


*Supplemental Figure 11. All tonic contraction TMS trials designated as "None". TMS onset at 0 s. "None" trials had no prominent EMG peak within the 10-ms search window. Peaks had to be above the $50^{th}$ percentile for peak prominence and above the $50^{th}$ percentile for peak width (for EMG peaks within the 1-second trial).*


Supplement for preprint deposited into *bioRxiv*. This version has not been peer reviewed.
*Ultrasound stimulation of the motor cortex during tonic muscle contraction.* (bioRxiv version 1)

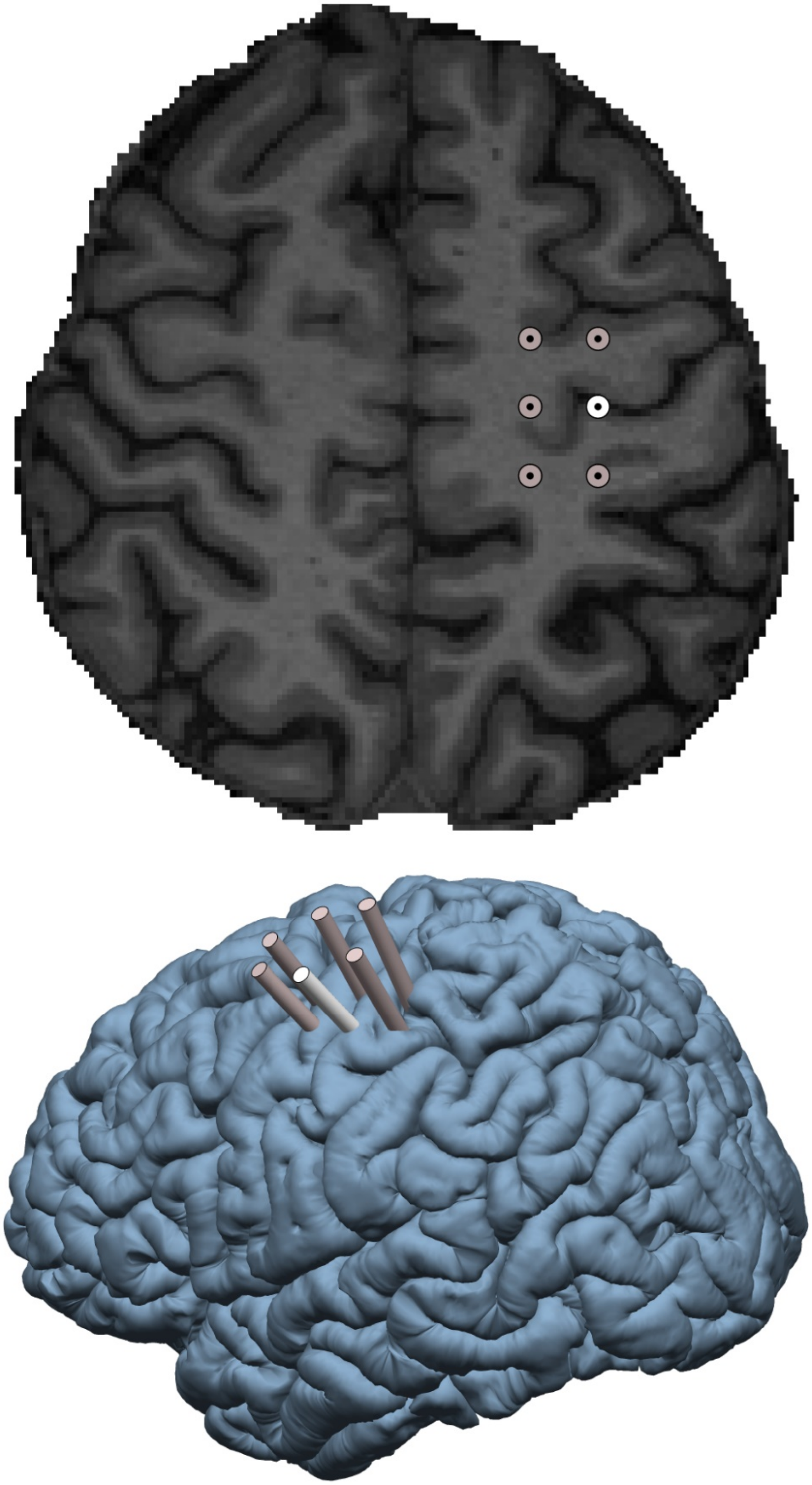

*Supplemental Figure 12. TMS search grid and trajectories. Illustration of the TMS search grid used in both 2D and 3D. The grid's origin (white) was placed at MNI coordinates that correspond to $M1_{hand}$ as based on a meta-analysis of fMRI motor experiments: x = −39, y = −24, z = 57* (Mayka et al., 2006). *The other five targets on the grid (grey) were in a 12 voxel-width grid (9.6 mm grid interval) around $M1_{hand}$ in subject space. See* EMG and NIBS Placement.

| Subject | FWHM [mm] | | | $Pressure_{Peak-to-Peak}$ [kPa] | | | Target |
|---|---|---|---|---|---|---|---|
| | Average | Dim. 1 | Dim. 2 | Anywhere | $M1_{hand}$ | Target | |
| sbj01 | 4.4 | 4.4 | 4.4 | 470 | 441 | 441 | $M1_{hand}$ |
| sbj01 | 4.6 | 4.8 | 4.4 | 441 | 37 | 399 | other |
| sbj01 | 4.9 | 5.0 | 4.8 | 406 | 19 | 378 | other |
| sbj07 | 4.5 | 4.4 | 4.6 | 428 | 386 | 386 | $M1_{hand}$ |
| sbj07 | 4.3 | 4.4 | 4.2 | 451 | 23 | 375 | other |
| sbj07 | 4.5 | 4.6 | 4.4 | 423 | 42 | 376 | other |
| sbj08 | 4.2 | 4.0 | 4.4 | 418 | 18 | 357 | other |
| sbj08 | 4.0 | 3.8 | 4.2 | 441 | 21 | 306 | other |
| sbj09 | 4.5 | 4.6 | 4.4 | 463 | 438 | 446 | $M1_{hand}$ |
| sbj09 | 4.9 | 5.0 | 4.8 | 434 | 41 | 414 | other |
| sbj09 | 4.9 | 4.6 | 5.2 | 414 | 13 | 343 | other |
| sbj10 | 4.4 | 4.0 | 4.8 | 430 | 15 | 280 | other |
| sbj10 | 4.2 | 4.2 | 4.2 | 456 | 34 | 326 | other |
| sbj10 | 3.9 | 3.6 | 4.2 | 439 | 13 | 316 | other |
| sbj11 | 4.6 | 4.6 | 4.6 | 374 | 361 | 361 | $M1_{hand}$ |
| sbj11 | 5.5 | 5.0 | 6.0 | 302 | 33 | 252 | other |
| sbj11 | 5.6 | 5.8 | 5.4 | 292 | 11 | 266 | other |
| sbj12 | 3.7 | 3.4 | 4.0 | 397 | 348 | 348 | $M1_{hand}$ |
| sbj12 | 4.4 | 4.4 | 4.4 | 362 | 25 | 268 | other |
| sbj12 | 4.6 | 4.8 | 4.4 | 367 | 10 | 274 | other |
| sbj13 | 4.4 | 4.2 | 4.6 | 430 | 395 | 395 | $M1_{hand}$ |
| sbj13 | 5.3 | 5.8 | 4.8 | 343 | 23 | 272 | other |
| sbj13 | 5.0 | 5.2 | 4.8 | 361 | 14 | 280 | other |
| sbj14 | 5.1 | 4.6 | 5.6 | 396 | 26 | 386 | other |
| sbj14 | 4.3 | 4.4 | 4.2 | 452 | 449 | 449 | $M1_{hand}$ |
| sbj14 | 5.0 | 5.0 | 5.0 | 412 | 18 | 372 | other |
| sbj16 | 5.2 | 4.4 | 6.0 | 386 | 314 | 314 | $M1_{hand}$ |
| sbj16 | 4.4 | 3.8 | 5.0 | 379 | 18 | 362 | other |
| sbj16 | 3.2 | 3.4 | 3.0 | 386 | 22 | 317 | other |

*Supplemental Figure 13. Table of simulated pressure values for each trajectory used with ultrasound. Data for both experiments included. Included are full width half maximum (FWHM) values [mm] of width of the ellipsoid focus of the focused ultrasound beam. The maximum pressures for three key locations for the simulation are also shown: the maximum pressure anywhere, at $M1_{hand}$, and at the target coordinate used to aim the trajectory. In some cases, the trajectory coordinate and the $M1_{hand}$ coordinate are the same ('Target' column).*


Supplement for preprint deposited into *bioRxiv*. This version has not been peer reviewed.
*Ultrasound stimulation of the motor cortex during tonic muscle contraction.* (bioRxiv version 1)

**Neurological screening questionnaire**

- Do you have any active medical, neurological, or psychiatric diagnosis (such as depression, schizophrenia, or bipolar disorder)?
- Do you have a pacemaker, deep brain stimulator, or other implanted electrical device (including intrauterine devices or braces)?
- Do you have, or have you ever had, a significant head injury or neurological disorder (such as a concussion or seizure disorder)?
- Do you have, or have you ever had, any seizures within the past six months?
- Do you have a family history of seizures?
- Do you have, or have you ever had, a history of alcohol or substance dependence?
- Do you have, or have you ever had, any cognitive impairments?
- Do you take any antidepressant medications (such as Prozac, Zoloft, or tricyclic antidepressants)?
- Do you take any antipsychotic medications?
- Do you take any antiviral medications?
- Do you take any amphetamines (such as Adderall)?
- Do you have a history of fainting?
- Do you have a history of migraines?
- Do you have a chronic pain disorder?
- Are you pregnant, or is there a chance you could become pregnant?
- Are you older than 50 years of age?

*Supplemental Figure 14. Full screening neurological health questionnaire used for recruitment. A 'Yes' to any question prevented inclusion in the study.*



Supplement for preprint deposited into *bioRxiv*. This version has not been peer reviewed.
*Ultrasound stimulation of the motor cortex during tonic muscle contraction.* (bioRxiv version 1)

#### Exposure Formula:

$$\sum_{traj=1}^{n} P_{traj} \times Time_{traj}$$

$\boldsymbol{n}$: Number of tUS trajectories

$\boldsymbol{P_{traj}}$: Pressure (est.) at $M1_{hand}$ voxel

$\boldsymbol{Time_{traj}}$: tUs-on time for that trajectory

#### k-Wave Parameters:

##### Medium Properties

| | **Density [kg/m3]** | **Speed of Sound [m/s]** | **Alpha Coefficient [dB/(MHz$^y$ cm)]** |
|---|---|---|---|
| **Skull** | 1732 | 2850 | 8.83 |
| **Brain** | 1546.3 | 1035 | 0.645946 |
| **Water** | 998 | 1482 | $6.7403 \times 10^{-5}$ |

Alpha Power (y): 1.43


Supplement for preprint deposited into *bioRxiv*. This version has not been peer reviewed.
*Ultrasound stimulation of the motor cortex during tonic muscle contraction.* (bioRxiv version 1)
